# LyGo: A platform for rapid screening of lytic polysaccharide monooxygenase production

**DOI:** 10.1101/2020.11.04.368555

**Authors:** Cristina Hernández-Rollán, Kristoffer B. Falkenberg, Maja Rennig, Andreas B. Bertelsen, Johan Ø. Ipsen, Søren Brander, Daniel O. Daley, Katja S. Johansen, Morten H. H. Nørholm

## Abstract

Environmentally friendly sources of energy and chemicals are essential constituents of a sustainable society. An important step towards this goal is the utilization of non-edible biomass as supply of building blocks for future biorefineries. Lytic polysaccharide monooxygenases (LPMOs) are enzymes that play a critical role in breaking the chemical bonds in the most abundant polymers found in recalcitrant biomass, such as cellulose and chitin. Predicting optimal strategies for producing LPMOs is often non-trivial, and methods allowing for screening several strategies simultaneously are therefore needed. Here, we present a standardized platform for cloning LPMOs. The platform allows users to combine gene fragments with different expression vectors in a simple 15-minute reaction, thus enabling rapid exploration of several gene contexts, hosts and expression strategies in parallel. The open-source LyGo platform is accompanied by easy-to-follow online protocols for both cloning and expression. As a demonstration, we utilize the LyGo platform to explore different strategies for expressing several different LPMOs in *Escherichia coli, Bacillus subtilis*, and *Komagataella phaffii*.

## Introduction

Transforming society from being dependent on fossil resources, to relying on sustainable resources is one of the most important challenges of the 21^st^ century. Overcoming this challenge is highly dependent on the displacement of petrochemicals with renewable biochemicals such as bioethanol, polylactic acid, and biosuccinic acid^1,2^. Currently, the majority of biorefineries in the EU utilize food crops, which gives rise to the food versus fuel debate and ultimately limits the implementation of these types of biorefineries^3,4^. This issue is mitigated by 2^nd^ generation biorefineries that rely on non-edible biomass as a resource, although depolymerization of this type of biomass constitutes a significant technical challenge^5,6^.

Lytic Polysaccharide Monooxygenases (LPMOs) are carbohydrate active enzymes that oxidize α and β-(1,4) glycosidic bonds in recalcitrant biopolymers, such as chitin and cellulose^7,8,9^. Due to the chemistry and architecture of the substrate binding surface, LPMOs generally act more readily on crystalline substrates compared to glycoside hydrolases (GHs)^7,10^. LPMO-induced cleavage weakens the structure of the biomass, and facilitates attack by other enzymes in a cascading mechanism^11,12^. Consequently, addition of LPMOs has been shown to increase the saccharification efficiency of commercially available cellulose cocktails^9,13^. Furthermore, LPMOs and LPMO-like proteins are involved in the virulence of plant^14^ and animal (including human) pathogens^15^. These factors make LPMOs highly interesting from both an industrial and academic perspective.

Heterologous expression of LPMOs is essential for characterization and industrial production of LPMOs. LPMOs have been produced in a range of organisms^16^. However, multiple expression strategies for the same enzymes are rarely explored or compared and previous efforts to standardize cloning into expression vectors in the LPMO field have been limited^17^. Ideally, a comprehensive strategy for heterologous expression of LPMOs enables simultaneous exploration of several strategies including e.g. different production hosts, gene sequence variants, signal peptides for secretion, localization within the hosts, solubility- and affinity tags, etc. Such a workflow would benefit from a simple standardized DNA editing approach and a diverse and accessible vector collection, which in addition would facilitate collaborations, automation, and data comparison.

Here, we describe an open-source platform, called “LyGo” (**Ly**tic polysaccharide monooxygenase **Go**lden gate cloning). The platform includes functionally validated expression vectors for three widely used protein production host: *Escherichia coli, Bacillus subtilis*, and *Komagataella phaffii* (formerly known as *Pichia pastoris*) and is accompanied by easy-to-follow online protocols for cloning and expression in each of the included organisms. It is our hope that the community will embrace, share, and expand the collection in the future. To demonstrate how the LyGo platform can be utilized, we explore a variety of expression strategies in the three production hosts. In all cases, the expression strategies show significant differences in performance, thus underlining why this type of synthetic biology resource is useful for improving production yields and for enabling sustainable biotech.

## Results

### LyGo cloning enables efficient and scarless assembly of expression vectors

Inspired by Golden Gate cloning^18^ and the Electra Vector System^®^ (ATUM, Newark, CA, USA), the LyGo platform utilizes the type IIS restriction endonuclease SapI to generate compatible DNA fragments. Because SapI cuts outside its recognition sequence, this allows for scarless assembly of DNA fragments containing LPMO-coding sequences (called “LyGo fragments”) into compatible expression vectors (called a “LyGo vectors”). SapI-treatment of a LyGo fragment creates three-nucleotide single stranded overhangs corresponding to the codon of the N-terminal histidine (essential for the activity of LPMOs^19^) and a stop codon in the 3’ end (Figure 1A). LyGo vectors contain a cloning cassette containing a *ccdB* counter selection marker^20^ flanked by SapI recognition sites (called a “LyGo cassette”). Upon SapI digestion, the LyGo cassette is released from a LyGo vector, leaving single stranded overhangs complementary to the digested LyGo fragments. This design ensures that any LPMO gene-of-interest, upon treatment with SapI and T4 ligase, can replace the cloning cassette without leaving cloning scars in the final vector. When a LyGo vector is combined with a LyGo fragment the SapI recognition sites are eliminated, and *in vitro* the correct assembly reaction is therefore favored over re-ligation of the LyGo cassette. Should any background vectors persist, the resulting transformants are selected against *in vivo* by the presence of the *ccdB* gene (Figure 1B, and Supplementary Figure S1A).

**Figure 1:**
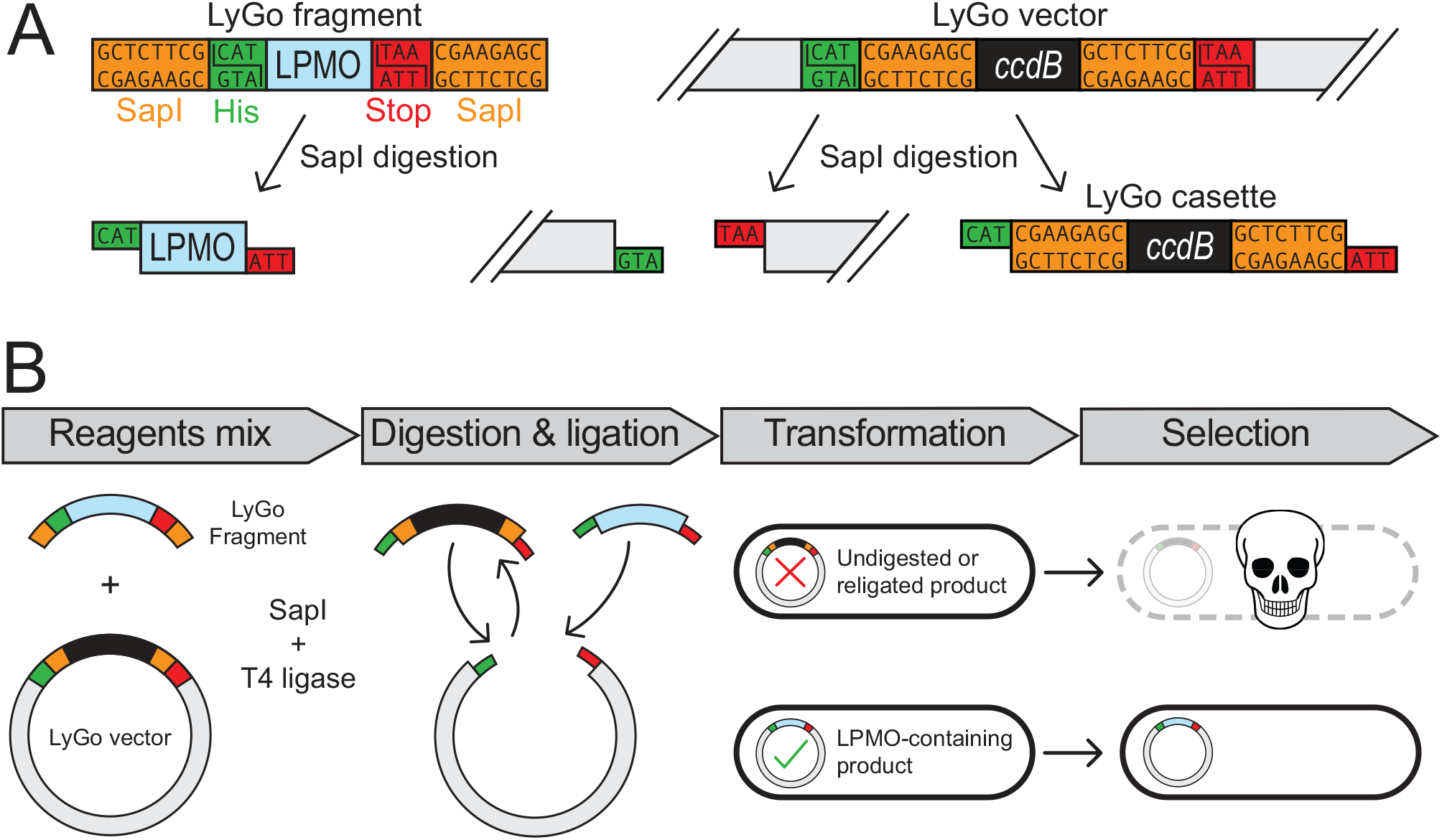
Schematic overview of LyGo cloning. **(A)** SapI recognition sites (orange), conserved histidine-(green), and stop codons (red) are shown in the LyGo-fragment, -cassette, and -vector before and after SapI digestion. **(B)** In the cloning procedure, LyGo fragment and vector are mixed with SapI restriction enzyme and T4 ligase, digested, and ligated in a 15-minute reaction, and subsequently transformed into a *ccdB* sensitive *E. coli* strain. This way, undigested or religated products are eliminated by *ccdB* counter selection.

Digestion and ligation are performed in a one-pot reaction at room temperature and can be completed in 15 minutes with high assembly efficiencies (Supplementary Figure S1B). The assembled DNA can be transformed directly into *E. coli*, following routine protocols^21^. The assembly is not dependent on DNA amplification or purification, which can be time-consuming, error-prone, and inefficient. A clear strength of this vector design is that any vector can be “LyGo-fied”, simply by removing unwanted SapI recognition sites and inserting the LyGo cassette. This allows users to utilize their favorite vectors and to contribute to the collection.

### Nomenclature for LyGo vectors

The current vector collection consists of 14 vectors in total (Table 1), and we established a simple abstraction and nomenclature, in order to easily recognize plasmids and translate their names into meaningful concepts. LyGo vectors are designated: “pLyGo” followed by two letters indicating the expression host (*Ec* for *E. coli, Bs* for *B. subtilis*, and *Kp* for *K. phaffii*), as well as a number that refers to information about the features of the plasmid. Any modules inserted by LyGo cloning of LyGo fragments are appended to the name separated by hyphens. For instance, pLyGo-*Ec*-2 is a LyGo vector for expression in *E. coli* that includes the T7 promoter and the signal peptide MalE^SP^ targeting LPMOs-of-interest to the periplasm. Cloning of a LyGo fragment encoding the LPMO LsAA9A into pLyGo-*Ec*-2, would be named pLyGo-*Ec*-2-LsAA9A.

**Table 1:** LyGo vectors constructed in this study. These vectors are available through plasmid repositories. TEV* denotes that the last codon of the TEV sequence was modified to a histidine codon.

| LyGo nomenclature | Classical nomenclature | Description |
| --- | --- | --- |
| *E. coli* vectors |  |  |
| pLyGo-*Ec*-1 | pET39b-Ub(his10)-TEV*-*ccdB* | Vector encoding Ubiquitin with an internal His10-tag and the LyGo cassette, Km<sup>R</sup> |
| pLyGo-*Ec*-2 | pET28a(+)-MalE<sup>SP</sup>-*ccdB* | Vector encoding the MalE signal peptide and the LyGo cassette, Km<sup>R</sup> |
| pLyGo-*Ec*-3 | pET28a(+)-OmpA<sup>SP</sup>-*ccdB* | Vector encoding the OmpA signal peptide and the LyGo cassette, Km<sup>R</sup> |
| pLyGo-*Ec*-4 | pET28a(+)-PhoA<sup>SP</sup>-*ccdB* | Vector encoding the PhoA signal peptide and the LyGo cassette, Km<sup>R</sup> |
| pLyGo-*Ec*-5 | pET28a(+)-DsbA<sup>SP</sup>-*ccdB* | Vector encoding the DsbA signal peptide and the LyGo cassette, Km<sup>R</sup> |
| pLyGo-*Ec*-6 | pET28a(+)-PelB<sup>SP</sup>-*ccdB* | Vector encoding the PelB signal peptide and the LyGo cassette, Km<sup>R</sup> |
| pLyGo-*Ec*-7 | pBAD42-PelB<sup>SP</sup>-*ccdB*-TEV-NB-C-IgAP | Vector encoding the C-IgAP surface expression anchor, nanobody, and the LyGo cassette, Km<sup>R</sup> |
| pLyGo-*Ec*-8 | pBAD42-Lpp<sup>SP</sup>-OmpA-TEV*-*ccdB* | Vector encoding the Lpp<sup>SP</sup>-OmpA surface expression anchor and the LyGo cassette, Km<sup>R</sup> |
| *B. subtilis* vectors |  |  |
| pLyGo-Bs-1 | pBS293C-amyE-P<sub>3P</sub>-AmyL<sup>SP</sup>-*ccdB* | Vector encoding the AmyL signal peptide and the LyGo cassette, Km<sup>R</sup> (*E. coli*), Cm<sup>R</sup> (*B. subtilis*) |
| pLyGo-Bs-2 | pBS293C-amyE-P<sub>3P</sub>-AmyQ<sup>SP</sup>-*ccdB* | Vector encoding the AmyQ signal peptide and the LyGo cassette, Km<sup>R</sup> (*E. coli*), Cm<sup>R</sup> (*B. subtilis*) |
| pLyGo-Bs-3 | pBS293C-amyE-P<sub>3P</sub>-AprE<sup>SP</sup>-*ccdB* | Vector encoding the AprE signal peptide and the LyGo cassette, Km<sup>R</sup> (*E. coli*), Cm<sup>R</sup> (*B. subtilis*) |
| pLyGo-Bs-4 | pBS293C-amyE-P<sub>3P</sub>-BatLPMO10<sup>SP</sup>-*ccdB* | Vector encoding the BatLPMO10 signal peptide and the LyGo cassette, Km<sup>R</sup> (*E. coli*), Cm<sup>R</sup> (*B. subtilis*) |
| ***K. phaffii* vectors** |  |  |
| pLyGo-Kp-1 | pPIC9K-α-MF<sup>SP</sup>-*ccdB* | Vector encoding the α-MF pre-sequence and the LyGo cassette, Amp<sup>R</sup> (*E. coli*), Km<sup>R</sup>, *HIS4* (*K. phaffii*) |
| pLyGo-Kp-2 | pPIC9K-Amy<sup>SP</sup>-*ccdB* | Vector encoding the α-amylase signal peptide and the LyGo cassette, Amp<sup>R</sup> (*E. coli*), Km<sup>R</sup>, *HIS4* (*K. phaffii*) |

### Exploring protein localization strategies in *E. coli* using LyGo

*E. coli* is an important model organism in molecular biology, and a popular expression host for bacterial LPMOs^16^. Proteins can be produced in different cellular compartments in *E. coli*, each strategy with its own advantages and disadvantages. In order to explore different approaches, we developed a range of vectors for producing LPMOs in three different cellular compartments of *E. coli*: The cytoplasm, the periplasm, and attached to the surface of the cell. The LyGo vectors were tested with four different LPMOs: *Thermobifida fusca* TfLPMO10A^22^, *Streptomyces coelicolor* ScLPMO10B^23^, BatLPMO10 from *Bacillus atrophaeus*^24^, and LsAA9A from *Lentinus similis*^25^. TfLPMO10A, ScLPMO10B, and BatLPMO10 were chosen because they previously were produced successfully in *E. coli*^22,23,24^. LsAA9A was chosen as an example of a fungal enzyme and because it had the added benefit of being compatible with a simple chromogenic assay (based on AZCL-HEC) available^26^. All sequences were codon optimized for *E. coli*, except for BatLPMO10 because the native sequence previously was expressed in *E. coli*^24^ at relatively high titers.

#### Cytoplasmic expression

We based a pLyGo-*Ec* cytoplasmic vector (pLyGo-*Ec*-1) on the pET39b backbone encoding an N-terminal ubiquitin solubility tag, a poly His-tag, and a TEV cleavage site^27^, which together has the potential to facilitate soluble expression, purification, and cleavage to release the N-terminal histidine (Supplementary Figure S2A).

The four different LPMO genes were inserted into pLyGo-*Ec*-1, transformed into *E. coli* BL21(DE3), and expression induced in late exponential phase by the addition of IPTG followed by incubation for 20 hours at 18 °C with shaking. Since TfLPMO10A, ScLPMO10B, and LsAA9A all contain disulfide bonds, we expressed them in parallel in the disulfide-bond enhancing strain *E. coli* SHuffle^28^. Using this setup, we observed that the different LPMOs were produced at titers ranging from approximately 100 to 800 mg/L (Figure 2B and Supplementary Figure S3), and that the disulfide containing LPMOs expressed considerably better in the SHuffle strain. However, the SHuffle strain grew to lower density, which reduced the difference in volumetric titers produced by the two strains.

**Figure 2:**
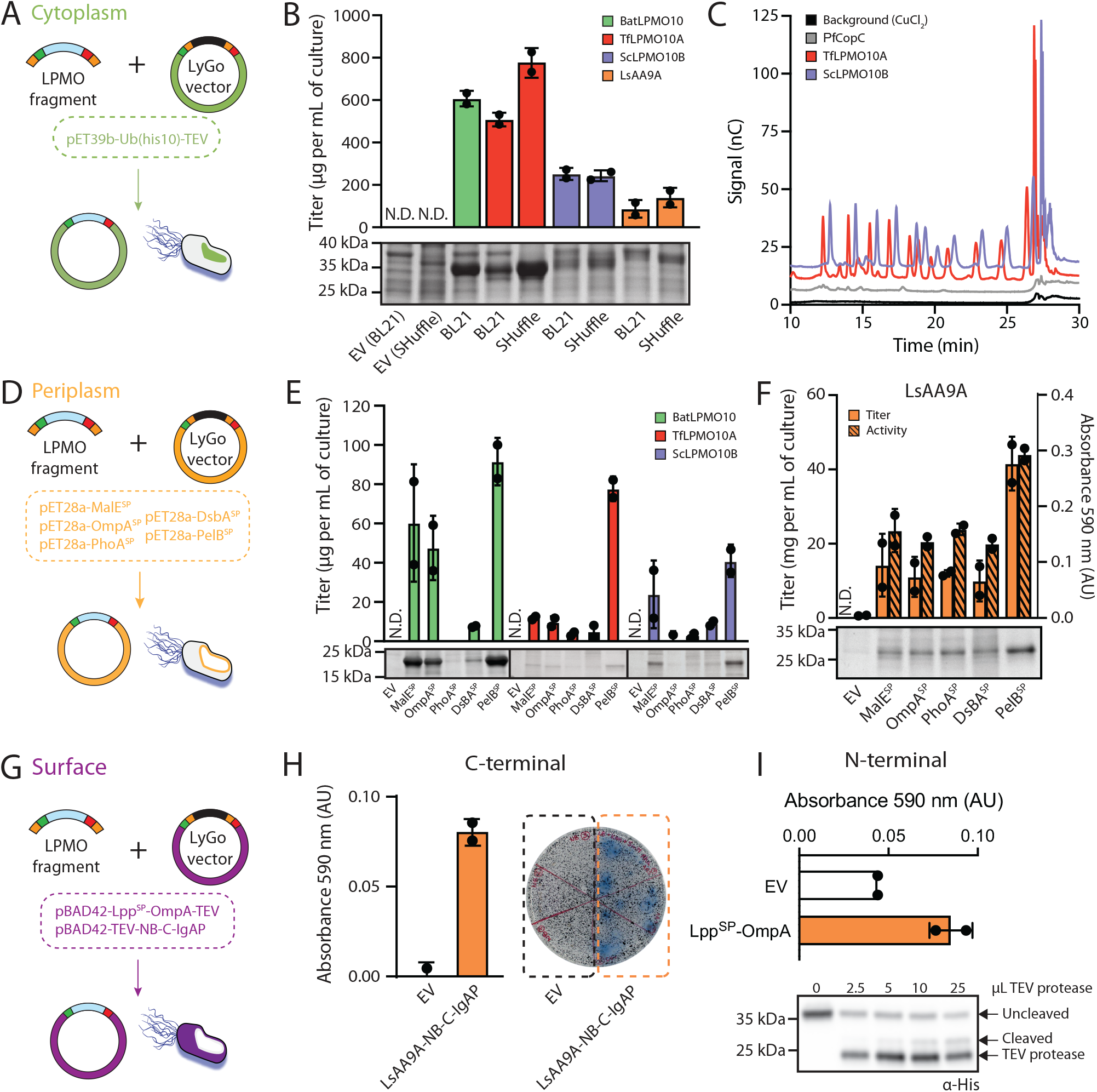
LPMO production in different compartments in *E. coli* using LyGo. EV = Empty vector, N.D. = Not determined. **(A)** Illustration of LyGo cloning for production of LPMOs in the cytoplasm of *E. coli*. **(B)** Results from cytoplasmic expression of four different LPMOs (see main text for further detail) in BL21(DE3) and the SHuffle strain. Protein production normalized by cell density was analyzed by densitometry of InstantBlue stained protein bands on an SDS-PAGE gel from biological duplicates (Supplementary Figure S3). A representative SDS-PAGE is shown in the lower panel. **(C)** Activities of TEV-cleaved LPMOs on PASC using HPLC-PAD, compared with the catalytically inactive control samples of PfCopC or 0.75 µM CuCl2. **(D)** Illustration of LyGo cloning for production of LPMOs in the periplasm with a range of different signal peptides. **(E)** Results from periplasmic expressions of BatLPMO10, TfLPMO10A, and ScLPMO10B. Production titers were quantified by densitometry of InstantBlue stained protein bands on an SDS-PAGE gel from biological duplicates (Supplementary Figure S5 to S7). A representative SDS-PAGE is shown in the bottom. **(F)** LsAA9A produced in the periplasm of *E. coli* with different signal peptides quantified by densitometry of InstantBlue stained protein bands on an SDS-PAGE gel from biological duplicates (Supplementary Figure S8), and activity of LsAA9A extracts quantified by the AZCL-HEC assay from biological duplicates. A representative SDS-PAGE from each construct is shown in the bottom. **(G)** Illustration of LyGo cloning for surface display of LPMOs with two different strategies. **(H)** Activity of the NB-C-IgAP surface displayed LsAA9A in liquid culture (left) and directly on LB agar plates using AZCL-HEC as a substrate (right). **(I)** LsAA9A expressed on the surface of *E. coli* using the N-terminal construct (pLyGo-*Ec*-8). Activity of the cleaved LsAA9A was analyzed by the AZCL-HEC assay (upper half). Western blot using an anti-His tag antibody for a His-tagged ScLPMO10B expressed on the surface of *E. coli* (bottom half). TEV protease was titrated to cleave the surface displayed protein.

It was possible to isolate all LPMOs by immobilized metal affinity chromatography (IMAC), but only TfLPMO10A and ScLPMO10B showed the expected fragmentation pattern after treatment with TEV protease (Supplementary Figure S4). The processed TfLPMO10A and ScLPMO10B were separated from the ubiquitin tag and the TEV protease by reverse IMAC purification (Supplementary Figure S4). Afterwards the enzymes were copper-loaded overnight and incubated with phosphoric acid swollen cellulose (PASC) and ascorbate for 25 hours and analyzed using high performance liquid chromatography with pulsed amperometric detection (HPLC-PAD). The assay showed a clear appearance of peaks corresponding to soluble oligosaccharides as observed previously for active LPMOs^29^. Furthermore, these peaks were absent when incubating the substrate with CuCl_2_ or with the non-catalytic Cu(II) chaperone CopC from *Pseudomonas fluorescens* SBW25 (PfCopC)^30,31^ (Figure 2C). This demonstrates cellulolytic activity from both LPMO-containing samples. Correct processing was confirmed by quantitative amino acid analysis (data not shown). Together, these results show that this expression system constitutes a viable strategy for producing active LPMOs in the cytoplasm of *E. coli*, although the efficiency of the crucial TEV cleavage step varies depending on the specific LPMO.

#### Periplasmic expression

Different signal peptides are known to influence translocation and folding kinetics of different proteins in an unpredictable manner^32,33^. Thus, screening of several different signal peptides is often key for optimizing expression and secretion into the periplasm. We based LyGo vectors for periplasmic expression (pLyGo-*Ec*-2 to pLyGo-*Ec*-6) on a pET28a(+) vector^34^ harboring sequences encoding the signal peptides MalE^SP^, OmpA^SP^, PhoA^SP^, and PelB^SP^ that target proteins to the periplasm post-translationally via the Sec-pathway. In addition, we included the DsbA^SP^ signal peptide as a representative of the co-translational SRP-pathway, as it previously has been shown to increase secretion of some proteins^35,36^. In an attempt to ensure high production titers, the translation initiation regions (TIRs) embedded in the coding sequence of the signal peptides, were replaced with the optimized versions described by Mirzadeh et al.^37^ (Supplementary Figure S2B).

The four LPMO constructs were transformed into BL21(DE3) and expression induced in late exponential phase by the addition of 1 mM IPTG and incubated for 20 hours at 18 °C with shaking, except for BatLPMO10 because pilot experiments showed improved expression at 30 °C. All the LPMOs were successfully expressed with titers of at least 40 mg/L culture (Figure 2E, 2F, and Supplementary Figure S5 to S8). The different signal peptides varied in performance for the different LPMOs, although PelB^SP^ showed the best overall performance. The periplasmic fractions collected from the expression of LsAA9A were subjected to AZCL-HEC activity assay (Figure 2F), which showed that all constructs produced functional versions of the enzyme with a good correlation between protein titer and enzymatic activity. This provides a demonstration that a fungal LPMO can be produced in a functional state in a bacterium using the LyGo platform.

#### Surface display and shaving

Surface display systems are useful tools for variant screening^38^ and whole cell catalysis^39^. Furthermore, controlled release from the membrane, as presented by Ahan et al.^40^, could be used as a method for selectively purifying the protein of interest. These concepts were materialized into two different designs: A C-terminal and an N-terminal fusion construct, which were both tested using LsAA9A. The pLyGo-*Ec* surface display vectors were based on a pBAD expression vector, which had previously been used for surface display^41^.

The C-terminal construct (pLyGo-*Ec*-7) uses the C-terminal translocation unit of the *Neisseria gonorrhoeae* autotransporter IgA protease (C-IgAP) and a single domain antibody (nanobody, NB), and should display active LPMOs (i.e. with the N-terminal histidine exposed) on the surface by fusing the NB to the C-terminus of the LPMO^41^. The C-terminal tag is obviously not compatible with the stop codon overhang in the standard LyGo design, so this overhang was changed to the first codon of a TEV site (glutamate, GAA) (Supplementary Figure S2C). The LsAA9A-encoding DNA sequence was cloned into this vector and transformed into BL21(DE3). Expression was induced with L-rhamnose in mid-exponential phase and the LPMO produced for 24 hours followed by functional AZCL-HEC assays in both liquid culture and agar plate format (Figure 2H). In both cases, activity was observed from the surface displayed LsAA9A, confirming that this design has the potential to be used for variant screening and whole-cell catalysis.

The N-terminal construct (pLyGo-*Ec*-8) uses a hybrid protein, consisting of the Lpp signal peptide and residues 66 to 180 of the outer membrane protein OmpA, and displays the LPMO as a fusion between the LPMO N-terminus and the OmpA C-terminus^41^. Furthermore, a sequence encoding a TEV site was inserted between the LyGo cassette and the Lpp^SP^-OmpA coding sequence, to allow for the release of the LPMO and the N-terminal histidine from the fusion protein upon treatment with TEV protease (Supplementary Figure S2C). LsAA9A was cloned into the construct, expressed (as described above), and released into the medium by addition of TEV protease. Following a centrifugation step, the supernatant was subjected to the AZCL-HEC assay. A small but significant increase in LPMO activity was observed indicating some successful cleavage and release of the surface displayed LsAA9A (Figure I top). However, due to the low signal observed in the assay, we speculate that TEV cleavage of LsAA9A is inefficient, as observed for LsAA9A TEV fusions expressed in the cytoplasm. To more convincingly demonstrate the surface “shaving”, we therefore decided to display and cleave a His-tagged version of ScLPMO10B with the same approach. Cleavage was induced by different concentrations of purified TEV protease (Figure 2I). Immunoblotting of the cleavage reaction mix with an anti-His tag antibody (α-His) showed that the fraction of anchored ScLPMO10B decreased while the fraction of free ScLPMO10B increased with increasing TEV concentrations. Together, this shows that LyGo vectors can facilitate functional surface display of LPMOs and that these can be released from the surface in a simplistic purification procedure.

### Exploring the effects of signal peptides and DNA sequence variants in *B. subtilis*

Bacteria in the *Bacillus* genus are industrially relevant, due to desirable traits such as high capacity for secreting proteins, ability to grow on cheap carbon sources, GRAS status, and robustness in industrial settings^42,43,44^. *B. subtilis* is widely used in both industry and academia, and the recent advances in standardizing genetic parts and strain libraries^45,46,47,48^, makes it an attractive protein production host.

LyGo plasmids for *B. subtilis* (pLyGo-*Bs*-1 to pLyGo-*Bs*-4) were constructed based on the integrative plasmid pBS293C-amyE from the *Bacillus* SEVA sibling collection^45^. Two SapI sites were removed from the backbone, and the LyGo cassette was inserted downstream of a triple promoter P_amyL_-P_amyQ_-P_cryIIIA-_cryIIIA_stab_^49^ and a predicted strong ribosome binding site (R0 from Guiziou et al.^48^). Like *E. coli*, signal peptide performance in *B. subtilis* is difficult to predict^50^, so we decided to generate four different plasmids with different signal peptides: AmyL^SP^ from *Bacillus licheniformis*, AmyQ^SP^ from *Bacillus amyloliquefaciens*, AprE^SP^ from *Bacillus clausii*, and BatLPMO10^SP^ from *Bacillus atrophaeus*. Initially, the plasmids were constructed similarly to the *E. coli* counterparts, but while assembling these we observed an unusual high number of frame-shift mutations in *ccdB*, despite the use of a *ccdB*-tolerant *E. coli* DB3.1. We hypothesized that this could be a result of toxicity caused by increased expression of *ccdB* from P_3P_ or other upstream sequences in the cloning host. In order to mitigate this, a stop codon was introduced in-frame with the signal peptide after the upstream SapI site, followed by a terminator from the BioBrick collection (BBa_B1002)^51^ (Supplementary Figure S2D). This updated design allowed for assembly of constructs with an intact *ccdB* gene in the LyGo cassette.

In addition to screening different signal peptides for, the pLyGo-*Bs* vectors were utilized to assess the expression of two different sequence variants of each of the sequences encoding BatLPMO10 and LsAA9A (Figure 3A): The native sequence and a sequence codon-optimized for *B. subtilis*. The resulting plasmids were transformed into the extracellular protease- and sporulation deficient *B. subtilis* KO7-S strain, thereby integrating the expression cassettes into the *amyE* locus by homologous recombination. The cells were grown in Cal18-2 media for 72 hours at 20 °C, the cells were harvested by centrifugation, and the supernatants analyzed by SDS-PAGE. BatLPMO10 produced at concentrations ranging from approximately 0.5 to 2 g/L in the supernatant (Figure 3B and Supplementary Figure S9). The signal peptides caused significant variation in expression levels, with the native BatLPMO10 signal peptide producing the highest amount of protein. The presence of LsAA9A in samples was barely detectable on SDS-PAGE gels stained by InstantBlue (data not shown). Thus, a His-tag was added to the sequences allowing for sensitive detection by Western blotting. Additionally, the activity of LsAA9A was assayed using the AZCL-HEC assay. The codon optimized variant was expressed at higher levels than the native sequence, roughly showing double the yield and activity compared to the native sequence (Figure 3C). The different signal peptides only yielded minor differences in activity, except for AprE^SP^ that performed poorly. Together, these results demonstrate that the pLyGo vectors can by utilized to produce and optimize secretion of functional LPMOs from both fungal and bacterial origin in *B. subtilis* and that industrially relevant (g/L) levels can be reached.

**Figure 3:**
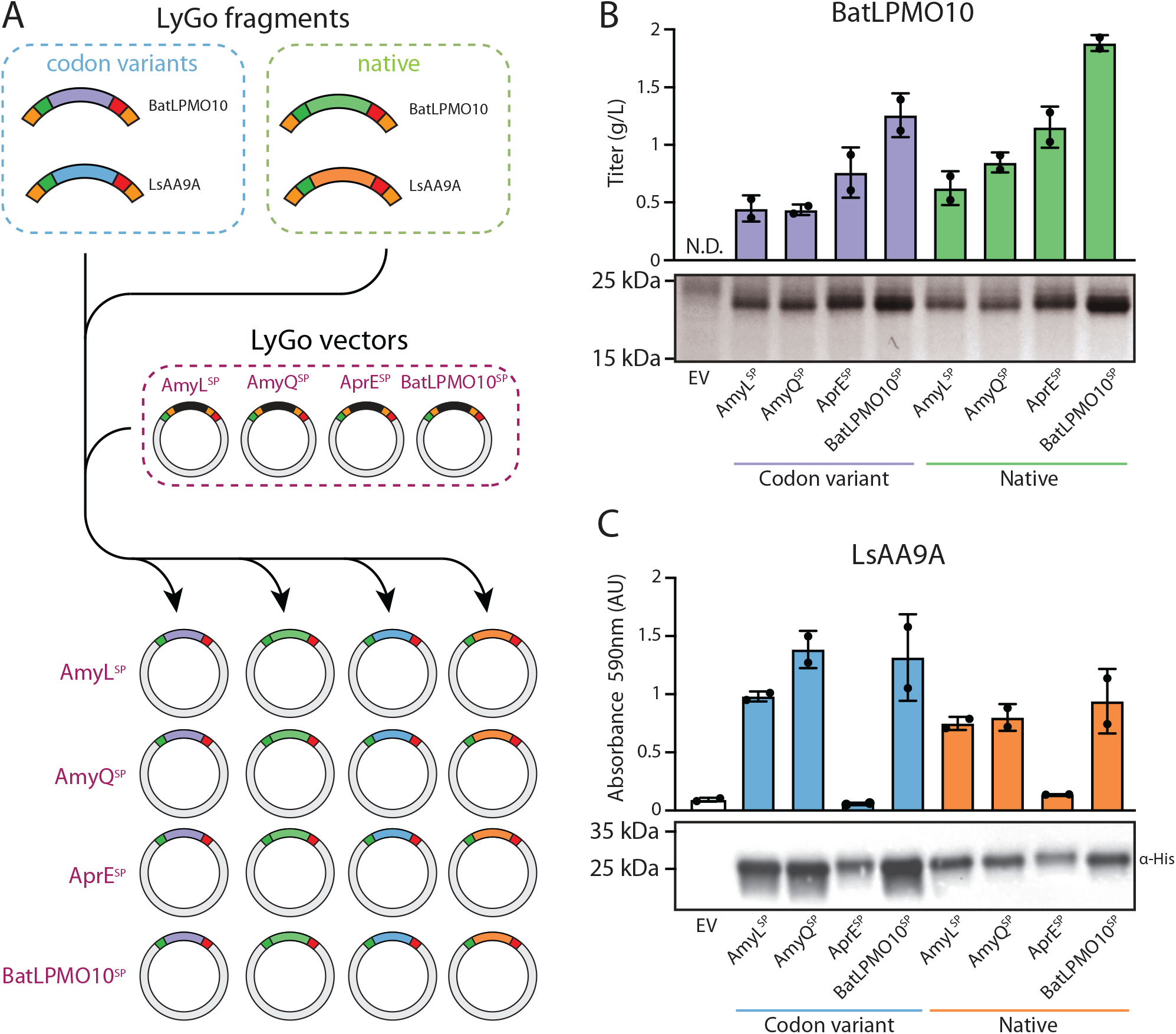
Exploration of sequence and signal peptide variants in *B. subtilis* using LyGo. EV = Empty vector, N.D. = Not determined. **(A)** Schematic overview of the combinatorial cloning strategy: Four different LyGo fragments encoding two sequence variants of two different LPMOs were cloned into four LyGo vectors encoding different signal peptides, resulting in a total of 16 expression vectors. **(B)** Analysis of expression of BatLPMO10 variants estimated by densitometry of InstantBlue stained protein bands on an SDS-PAGE gel from two biological replicates (Supplementary Figure S9). A representative gel picture is shown below. **(C)** Activity of LsAA9A was quantified by AZCL-HEC assay from two biological replicates. A representative Western-blot of each variant is shown below.

### Exploring the effects of signal peptides and sequence variants in *K. phaffii*

The methylotrophic yeast *K. phaffii* is the most frequently used yeast species for production of recombinant proteins^52^. This is due to its ability to grow to high cell density, to express recombinant genes in a tightly controlled manner, and to efficiently secrete proteins^52,53,54,55^. Despite its biotechnological importance and wide use in industry, relatively few genetic tools are readily available. However, *K. phaffii* has proven to be a promising production host for carbohydrate active enzymes like LPMOs^16,56,57,58,59^ and could be a valuable addition to the LyGo platform.

LyGo vectors for *K. phaffii* (pLyGo-*Kp*-1 and pLyGo-*Kp*-2) were created based on the pPIC9K plasmid hosting the methanol-inducible P_AOX1_ promoter. A multiple cloning site downstream of the α-mating factor pre-sequence (α-MF^SP^) from *Saccharomyces cerevisiae* was replaced by the LyGo cassette and a SapI site was removed from the backbone by site-directed mutagenesis. A second vector (pLyGo-*Kp*-2) was designed, where α-MF^SP^ was replaced with the α-amylase signal peptide (Amy^SP^) from *Aspergillus niger* (PichiaPink™ Secretion Signal Kit, Thermo Fisher) (Supplementary Figure 2E).

Two sequence variants of the gene encoding LsAA9A were cloned into the two pLyGo-*Kp* vector*s*: the native sequence and a sequence variant codon optimized for *K. phaffii*. The resulting plasmids were linearized and integrated into *K. phaffii* GS115^60^. This type of integration results in two distinct phenotypes: Mut^+^ and Mut^S^, which have shown to impact recombinant protein production^61^. In total, this leads to eight different strain variants which were tested for activity on the AZCL-HEC substrate (Figure 4A). Protein production was performed over 4 days at 28 °C with daily addition of methanol, before the cells were harvested and the supernatants were analyzed by SDS-PAGE. LsAA9A was produced at concentrations up to approximately 0.3 g/L (Figure 4B). The expression was largely unaffected by codon variation and strain genotype, and a trend was that the α-MF^SP^ signal peptide produced at higher levels than Amy^SP^. Protein amounts were found to correlate poorly with the cellulolytic activity measured by the AZCL-HEC assay (Figure 4B): despite the relatively low production titers from the Mut^+^ strain harboring LsAA9A with the Amy^SP^, the sample showed the highest overall activity. In summary, the *K. phaffii* LyGo platform expands the possibilities of the LyGo platform to include screening of expression strategies in an eukaryotic host.

**Figure 4:**
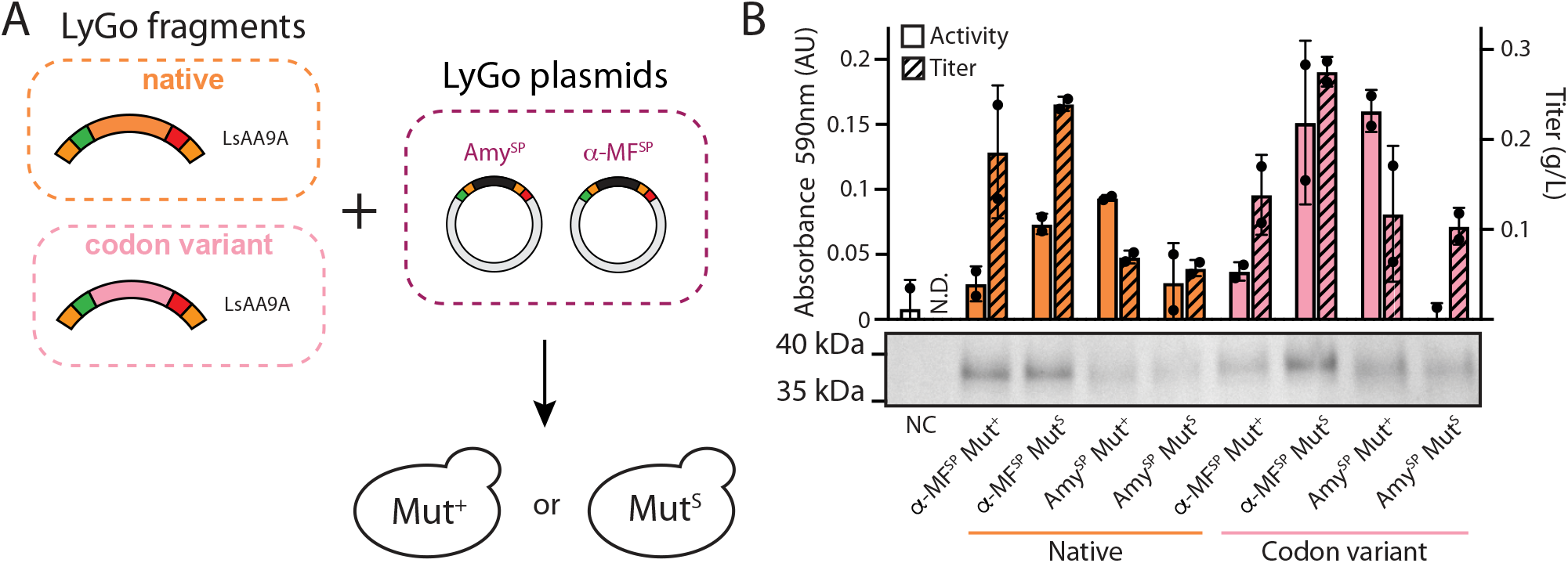
Production and activity of LsAA9 with sequence and signal peptide variants in *K. phaffii* using LyGo. EV = Empty vector, N.D. = Not determined. **(A)** Schematic overview of the combinatorial cloning strategy: LyGo fragments encoding the native sequence and a codon variant of LsAA9A were cloned into two LyGo vectors encoding different signal peptides and the resulting constructs integrated in *K. phaffii* resulting in Mut^+^ and Mut^S^ phenotypes. **(B)** Expression was analyzed by densitometry of InstantBlue stained protein bands on an SDS-PAGE gel, and activity quantified by AZCL-HEC assay from two biological replicates (Supplementary Figure S10). A representative gel picture is shown in the lower panel.

## Discussion

Progress in the LPMO field ultimately depends on the availability and quality of the enzyme samples used in different laboratories. Thus, failed or low titer expression is a limiting factor for advancement in the field. Typically, most research groups will work with a limited number of expression strategies, and may sometimes fail in expressing a specific gene or accept low-titer samples. The LyGo platform provides easy access to a range of expression strategies that can be explored in parallel in order to increase the likelihood of high-titer and high-quality expression. Furthermore, standardization of the cloning and expression workflows may improve sample uniformity and comparability of results across different laboratories. In order to facilitate the use of LyGo, the vectors are made available through plasmid repositories such as Addgene, and detailed protocols are available at the online platform protocols.io (Supplementary figure S11).

In this work we demonstrate that LyGo can be used for identifying an optimal expression context for different LPMOs in three different organisms. For *E. coli*, we developed expression vectors for cytoplasmic, periplasmic, and surface expression. Cytoplasmic expression is the most common strategy for the production of heterologous proteins in *E. coli*^62^, and is often advantageous over other expression strategies as it mitigates the need for screening signal peptides and has the potential to result in product yields exceeding 50% of the total cellular protein amounts^63^. Furthermore, cytoplasmic expression supports the use of several genetic tools such as co-expression of chaperones^64^ and folding sensors^65^, as demonstrated in this work using the *E. coli* SHuffle strain. However, cytoplasmic expression is not the most obvious strategy for production of LPMOs for two reasons: (1) a functional LPMO requires a free N-terminal histidine residue^19^, which is made impossible by the start-codon-encoded methionine in standard cytoplasmic expression strategies. (2) LPMOs frequently contain disulfide bonds^66^, which are not readily formed in the cytoplasm of *E. coli* due to the reducing environment^67^.

We mitigated the first issue by inserting an N-terminal tag, which can be removed by TEV cleavage leaving an N-terminal histidine residue and demonstrated how this results in an untagged and functionally active LPMO. Furthermore, an N-terminal tag has several potential advantages: It can enhance expression and solubility of the fusion protein, and it can it can allow for purification by IMAC purification for both initial separation prior to TEV cleavage and for subsequent removal of the tag and the TEV protease. Finally, it could quench unwanted oxidations within the production host, which may lead to production of reactive oxygen species^68^. Similar methods have been used to express a range of LPMOs^69,70,71,72^, although none of these methods take advantage of a solubility tag. Unfortunately, we did not observe any TEV cleavage for BatLPMO10 and LsAA9A, potentially due to steric hindrance at the Ubiquitin/LPMO interface. Thus, this strategy is not a one-size-fits-all strategy for LPMOs. On the other hand, we were able to express and effectively TEV-cleave TfLPMO10A and ScLPMO10A at significantly increased titers compared to the same LPMOs expressed by the periplasmic expression strategy, demonstrating that this is an attractive strategy for preparing high titer samples of LPMOs.

In order to facilitate formation of disulfides, we expressed LPMOs in the *E. coli* SHuffle strain, which enhances the formation of disulfide bonds in the cytoplasm^28^. This significantly improved the overall titer of TfLPMO10A, but had little effect on the titers of ScLPMO10B and LsAA9A due to the poorer growth of the Shuffle strain.

Periplasmic expression is a popular strategy for expressing LPMOs in *E. coli*^16^, as it naturally provides the means for N-terminal processing and disulfide bond formation. However, secretion has previously been shown to constitute a bottleneck resulting in low yields^73^. Furthermore, this strategy often requires laborious screening of different signal peptides in order to optimize performance^37^. In order to simplify such a workflow, we provide vectors for screening several signal peptides in parallel, thereby reducing the time and effort required for this type of optimization. Of the tested signal peptides, we found PelB^SP^ to consistently perform the best. It is unclear whether this is a prevalent property of PelB^SP^, or an anecdotal observation due to the small number of tested LPMOs.

Production of proteins on the surface generally suffers from similar bottlenecks as periplasmic expression. A previous study has shown that different signal peptides affect the display efficiency of the N-terminal surface display construct^41^. In the specific case of LPMOs, while the N-terminal surface display construct ensures proper processing, the C-terminal surface display construct requires cleavage by TEV protease for the LPMO to be active. Unfortunately, in line with our observations from cytoplasmic production, TEV cleavage of LsAA9A was inefficient, providing a cautionary tale that the C-terminal surface display construct might not be suitable for all LPMOs. While TEV protease can be produced at low cost in *E. coli*, simultaneous production of a surface displayed TEV protease might present an attractive extension of the system.

For *B. subtilis* and *K. phaffii* we both explored the effect of signal peptides and sequence variants. As observed many times previously, the results from *B. subtilis* expression show that both signal peptides and codon optimization efforts influence expression in a protein-dependent manner. This underlines the usefulness of a simple and efficient platform for exploration of expression strategies and importantly our pLyGo-*Bs* vectors showed capabilities to express functional LPMOs at industrially relevant titers.

The *K. phaffii* LPMO production from pLyGo vectors also showed variations in performance between signal peptides, codon variants, and phenotypes, but with complex interplay between these factors. Furthermore, the production titers and measured activities were found to correlate poorly. One explanation could be the promiscuity in the processing of the α-MF^SP^ signal peptide^74,75^, leading to inefficient release of the N-terminal histidine, but it is also possible that the high concentration of copper sulphate used in the AZCL-based assay was suboptimal for a proper quantitative readout.

In addition to expressing LPMOs at different titers, different hosts can also give rise to different protein modifications (such as methylation and glycosylation^76^). The choice of organism(s) can therefore be a strategy to optimize protein stability, solubility or activity^77^ - or as a means to study the importance of these modifications. Some fungal LPMOs have been found to be selectively methylated at the N-terminal histidine^9^. This modification has been suggested to protect the enzymes from oxidative inactivation^76^, although this is still actively being investigated. The LyGo platform currently supports the non-glycosylating and non-methylating hosts *E. coli* and *B. subtilis*, and the glycosylating and non-methylating host *K. phaffii*. A methylating host would therefore be a valuable future addition to the platform, as LyGo then would cover the most important modifications for LPMOs.

It is our hope that the community will adopt and expand the LyGo platform with vectors and protocols for additional expression strategies. For this, careful design of the LyGo vectors is necessary. During construction of the pLyGo vectors, we learned that building constructs with *ccdB* is sensitive to the context of the LyGo cassette – possibly due to increased expression of *ccdB* facilitated by upstream sequences. In one case, this was solved by inserting an in-frame stop codon and a terminator in the pLyGo-*Bs* vectors (Supplementary Figure S2). Future additions to this or similar vector collections should consider this potential issue early in the design phase.

## Materials and Methods

### Strains

The strains used in this study are listed in Supplementary Table 1. *E. coli* DB3.1 (Invitrogen, Carlsbad, CA, USA) was used to construct and propagate *ccdB*-containing plasmids. *E. coli* NEB5α (New England Biolabs, Ipswich, MA, USA) was used to construct and propagate of LPMO-containing constructs. Expression experiments were performed with *E. coli* BL21(DE3) (Novagen, Merck KGaA, Darmstadt, Germany), *B. subtilis* KO7-S (The Bacillus Genetic Stock Center, Columbus, OH, USA), and *K. phaffii* GS115 (Thermo Fisher Scientific, Waltham, MA, USA). Bacterial cells were routinely cultivated in lysogeny broth (LB) at 37 °C with 250 RPM of shaking or on LB agar plates at 37 °C. When appropriate, the media was supplemented with kanamycin (50 µg/mL) for *E. coli* and chloramphenicol (5 µg/mL) for *B. subtilis*. Competent *E. coli* cells were obtained from (Invitrogen, California, United States) and transformed following standard protocols. Competent *B. subtilis* cells were obtained as described elsewhere^78^, although without adding histidine to the SM1 and SM2 media and with at least 2 hours of recovery. Competent *K. phaffii* cells were obtained using the Pichia EasyComp Kit (Invitrogen, Thermo Fisher Scientific, Waltham, MA, USA).

### Vector construction

DNA manipulations were performed using uracil excision cloning as described elsewhere^79^. The plasmids used and constructed in this work are listed in Table 1 and Supplementary Table 2. PCRs were performed using Phusion U Hot Start polymerase (Thermo Fisher Scientific, Waltham, MA, USA), according to manufacturer’s instructions. Primers were ordered from Integrated DNA technologies (IDT, Coralville, IA, USA). The primers used in this work are listed in Supplementary Table 3. Plasmids sequences were confirmed by sequencing (Eurofins MWG operon, Germany).

### LyGo cloning

Inserts were either synthesized as gBlocks (IDT, Coralville, IA, USA) or Strings (Thermo Fisher Scientific, Waltham, MA, USA), or amplified by PCR using Phusion Hot Start II DNA polymerase (Thermo Fisher Scientific, Waltham, MA, USA), according to manufacturer’s instructions. The sequence of the codon variants were obtained using IDT’s Codon Optimization Tool or Thermo Fisher’s GeneOptimizer Tool^80^ LyGo cloning was performed by mixing vector and insert in molar ratios between 1:3 to 1:6, together with 0.5 µL FastDigest LguI (SapI) (Thermo Fisher Scientific, Waltham, MA, USA), 1 µL FD buffer (Thermo Fisher Scientific, Waltham, MA, USA), 1 µL T4 DNA Ligase (Thermo Fisher Scientific, Waltham, MA, USA), and 1 µL T4 DNA Ligase buffer (Thermo Fisher Scientific, Waltham, MA, USA). The reaction volume was adjusted to 10 µL using MilliQ water, incubated at room temperature for at least 15 minutes, and subsequently used for transformation.

### Expression in *E. coli*

The expression of LPMOs in the periplasm and cytoplasm of *E. coli* performed similarly as in Hemsworth et al.,^81^ a single colony of the desired strain was inoculated in 50 or 25 mL of LB medium supplemented with kanamycin and grown O/N at 37 °C with 250 RPM. The O/N culture was back-diluted 1:100 in 50 mL of fresh LB medium with kanamycin and incubated at 37 °C with 250 RPM. When an OD_600_ value between 0.5 and 0.6 was reached, the cultures were transferred to an 18 °C incubator at 180 RPM (unless otherwise stated) and grown until an OD_600_ value between 0.8 and 1. At this point, the expression was induced with 1 mM IPTG and the cultures were grown for 20 hours at 18 °C incubator at 180 RPM (unless otherwise stated). Cultures were normalized to equal OD_600_ values corresponding to 25 OD units (ODU) using LB medium, spun down at 8000 g for 20 minutes at 4 °C, and the supernatant was discarded.

For surface expression the cultures were treated similarly, although the culture was induced with 5 mM L-rhamnose once an OD_600_ of 0.5 was reached. The culture was incubated at 30 °C at 250 RPM for 20 hours, and the cells directly plated on LB agar plates containing the AZCL-HEC substrate or harvested at 4000 g for 5 minutes, washed twice and resuspended in 10 mM Tris-HCl pH 7.5 (100 μL per ODU) for activity assay in liquid culture. For cleavage of the surface displayed protein between 1 and 25 μL of purified TEV protease was added and the reaction incubated at room temperature overnight. Subsequently, cells were pelleted at 5000 g for 10 minutes and the supernatant was collected for analysis by SDS PAGE, western blot or activity assay.

### Whole-cell lysis

For the screening of production conditions, extraction of the soluble fraction of the whole-cell lysate was done by chemical lysis using Cellytic™ B (Sigma-Aldrich, Saint Louis, MO, USA). Every ODU of cells was lysed using 16 µl of Cellytic™ B in 50 mM tris pH 8, 10mM imidazole, 150 mM NaCl buffer (supplemented with benzonase and lysozyme according to batch size). Following lysis, the insoluble fraction was pelleted by centrifugation at 25,000 g for 25 min at 4 °C and the supernatant stored at 4 °C until use. For the SDS-PAGE gels of whole-cell lysates, 0.046 ODU was loaded in each lane. For expressions in 500 mL culture volume for production of TfLPMO10A and ScLPMO10B for activity testing, cells were lysed by freeze-thawing. The pelleted cultures were frozen at –80 °C for an hour before the cells were thawed at RT and resuspended in 50 mL lysis buffer (50 mM tris pH 8, 10mM imidazole, 150 mM NaCl buffer). The lysis was now achieved by two cycles of at least 1 hour at −80 °C followed by thawing at room temperature. Following lysis, the insoluble fraction was pelleted by centrifugation at 15,000 g for 25 min at 4 °C and the supernatant stored at 4 °C until use.

### Periplasmic protein extraction

The periplasmic fraction was isolated following a modified version of an existing protocol^82^ by adding 12 µl of TSE buffer (200mM Tris-HCl pH 8, 500mM sucrose, 1mM EDTA) per ODU collected. The pellet was resuspended by mild pipetting, and the mixture was incubated at room temperature for 10 minutes. Afterwards, the cultures were subjected to a cold shock by adding 12 µl of ice-cold water per ODU. Finally, the mixture was subjected to a centrifugation step of 8000 g for 20 minutes at 4 °C and the pellet was discarded. Supernatants containing the periplasmic fraction were kept at 4 °C for enzymatic assays. Following this modified protocol, we were able to obtain substantially cleaner and purer periplasmic fractions in which lysozyme was not added to the buffer mixture (Supplementary Figure S12). 0.16 ODU was loaded in each lane of the gels for SDS-PAGE analysis.

### IMAC and reverse IMAC purification

Initial small test purifications were performed using Ni-NTA Spin Column for His-Tagged proteins (Qiagen, Hilden, Germany) following the protocol for 6xHis-Tagged Proteins under Native Conditions from *E. coli* Cell Lysates. Large batch purifications were performed using Ni-NTA Superflow (Qiagen, Hilden, Germany). A volume of lysate equivalent to 10-20 mg of the target protein was mixed with lysis buffer (50 mM Tris-HCl pH 8, 10 mM imidazole, 150 mM NaCl) to a final volume of 16 mL. This mixture was added to 2 mL of equilibrated Ni-NTA Superflow beads and incubated at 4 °C for 25 min with slow stirring. The flow-through was discarded, and the sample was washed twice with 15 mL wash buffer (50 mM Tris-HCl pH 8, 20 mM imidazole, 150 mM NaCl). Finally, the target fusion protein was eluted from the column with 5 mL elution buffer (50 mM Tris-HCl pH 8, 300 mM imidazole, 150 mM NaCl). The total volume of lysate from where the protein was purified was 9, 12, and 48 mL for PfCopC, TfLPMO10A, and ScLPMO10B, respectively. After TEV-cleavage, EDTA was removed by gel filtration on a 220 mL Sephadex-G25 (Cytiva, São Paulo, Brazil) column setup to an ÄKTA pure chromatography system. 30 mL of TEV treated sample was loaded on the column, followed by elution with dialysis buffer (50 mM tris pH 8, 200 mM NaCl). The protein content of the sample was collected between 70 to 110 mL retention volume. Afterwards, the target protein was separated from the His-tagged ubiquitin and TEV protease by running the sample over 5 mL equilibrated Ni-NTA Superflow beads and collecting the flow-through. Effective removal of the Ub(his10)-tag and TEV protease was ensured by SDS-PAGE (Supplementary Figure S4), and confirmed by amino acid analysis as described by Barkholt et al.^83^ Subsequently, the samples were concentrated using Amicon ultra spin filters (Sigma-Aldrich, Saint Louis, MO, USA) (10.000 MWC for TFLPMO10A and ScLPMO10B and 5.000 MWC for PfCopC).

### Production and purification of TEV protease

pTEVprotease was transformed into *E. coli* BL21(DE3) cells. A fresh colony was inoculated into LB medium supplemented with ampicillin at 37 °C with 250 RPM shaking overnight. The next day, the culture was diluted to an OD_600_ of 0.1 and incubated at 37 °C with 250 RPM shaking until an OD_600_ of 0.9 at which point the cells were induced with 1 mM IPTG and incubated overnight at 37 °C. The next day, cultures were collected by centrifugation and the pellets resuspended in lysis buffer (50 mM Tris-HCl, 200 mM NaCl, 10 mM imidazole, pH 7.8). The resuspended pellets were subjected to sonication in order to obtain the protein fraction (10 intervals of sonication for 10 seconds at the time with 30 seconds pause in between) while maintaining the cell lysates on ice. Afterwards, the sonicated slurry was centrifuged for 1 hour at 15,000 g at 4 °C and the supernatant containing the protein fraction was kept at 4 °C until purification. TEV protease purification was performed by IMAC purification. The resulted in purified TEV protease fractions which were loaded on SDS-PAGE to identify the fractions. The TEV protease was buffer exchanged (50 mM Tris pH 8, 150 mM NaCl) and concentrated in Amicon ultra spin filters (Sigma-Aldrich, Saint Louis, MO, USA) (10.000 MWC) to an ABS_280_ ≥ 3.0, before activity was confirmed and the sample stored as aliquots with 50% glycerol at −80 °C.

### Cleavage with TEV protease

Immediately before the cleavage, the TEV protease was reduced with 10 mM of 1,4-dithiothreitol (DTT, Sigma-Aldrich, Saint Louis, MO, USA) at room temperature for 30 minutes. For the cytoplasmically produced proteins, TEV protease was mixed with the target fusion protein in a 1:10 ratio based on Abs_280._ Before adding the protease to the reaction mixture, 2 mM EDTA was added to the reaction. The reaction was incubated at 16 °C overnight. For the proteins expressed on the surface 5 µL of TEV protease were mixed with 1 ODU of cell suspension (unless otherwise stated). The reaction was incubated at RT overnight.

### HPLC-PAD-based cellulose degradation assay

Proteins were preloaded at 4 °C overnight with CuCl2 at 1:0.75 stoichiometry. The assay was performed in 200 µL samples containing 50 mM citrate phosphate buffer pH 7.25, 0.25 % phosphoric acid swollen cellulose (PASC), 1 mM ascorbic acid and 0.75 µM copper-loaded enzyme. The samples were prepared in microtiter plates in technical triplicates. Samples where incubated for 25 hours in a thermomixer at 50 °C with 750 RPM shaking. Samples where filtered and analyzed by HPLC-PAD as described by Westereng et al.^84^.

### AZCL-HEC-based cellulose degradation assay

Activity of LsAA9A samples was measured by mixing a reaction solution consisting of 1 mg/mL AZCL-HEC substrate (Megazyme, County Wicklow, Bray, Ireland), 2 mM Ascorbic acid (Sigma-Aldrich, Saint Louis, MO, USA), 200 µM Copper sulphate (Sigma-Aldrich, Saint Louis, MO, USA), and adjusted to the desired volume with 100 mM Sodium acetate (pH 5) (Sigma-Aldrich, Saint Louis, MO, USA). 100 µL sample was mixed with 400 µL reaction mix and incubated at 50 °C with 1500 RPM shaking until a blue color had developed in the samples with LsAA9A. The reactions were centrifuged at 17000 g for 5 min and the absorbance of the supernatant was measured at 590 nm. Controls with the buffer or media the enzymes were suspended in were included, and the absorbance values of these samples were subtracted from the other measurements. For plate experiments, a 2x LB agar solution was mixed in a 1:1 ratio with a 2x reaction buffer (final concentration as described above) and poured very thinly to assure that the AZCL-HEC substrate was accessible from the surface.

### Expression in *B. subtilis*

*B. subtilis* transformants were streaked on LB agar plates supplemented with 1% starch, incubated at 37 °C for 24 hours, and a strain which had not produced a visible halo (indicating a correct double crossover event) was selected. A single colony was inoculated in 3 mL of LB medium supplemented with chloramphenicol and grown O/N at 37 °C with 250 RPM shaking in an Axygen 24-deepwell plate (Corning Life Science, Corning, New York, USA). The O/N culture was back-diluted 1:300 in 3 mL fresh Cal18-2 medium^85^ (Glucidex 12 was exchanged for Maltodextrin DE 13-17 (Sigma-Aldrich, Saint Louis, MO, USA) supplemented with chloramphenicol also in an Axygen 24-deep well plate. The culture was grown for 72 hours at 20 °C with 250 RPM shaking. The cells were harvested by centrifugation at 6000 g for 5 minutes at 4 °C, and the pellet was discarded. The samples were stored at 4 °C before use. 10 µL were loaded on gels for SDS page and western blotting.

### Expression in *K. phaffii*

A single colony of the desired strain was inoculated in 25 mL pH 6 buffered Glycerol-complex (BMGY) medium (*Pichia* Expression Kit, Life Technologies, Carlsbad, CA, USA) and grown at 28 °C with 250 RPM shaking until saturation. The preculture was then diluted 1:40 into 100 mL of pH 6 buffered BMGY media and grown O/N at 28 °C with 250 RPM shaking. Cells were pelleted, and resuspended in Buffered Methanol-complex (BMMY) medium (*Pichia* Expression Kit, Life Technologies, Carlsbad, CA, USA) to a final OD_600_ of 50. This suspension was back-diluted 1:50 in a 500 mL baffled shake flask containing 100 mL BMMY medium. Expression was performed for 4 days at 28 °C with 250 RPM shaking. Induction was maintained by addition of 1% methanol to the cultures once a day. The cells were harvested by centrifugation at 4000 g for 20 minutes at 4 °C and the pellets were discarded. The samples were stored at 4 °C before use. 10 µL were loaded on gels for SDS page.

### SDS-PAGE analysis

Enzyme samples were mixed 1:1 with sample buffer (8 M urea, 0,0105 % (w/v) bromophenol blue, 5 mM EDTA, 100mM Tris-HCl pH 6.8, 4 % (w/v) SDS, and 25% (v/v) glycerol), and heated to 98 °C for 10 minutes. Specific volumes of samples (described previously) were loaded on a 4−20 % Mini-PROTEAN-TGX gel (BioRad, Hercules, CA, USA) and run at 180 V for 45 min. Gels were stained for at least 4 hours using InstantBlue Protein Stain (Expedeon Inc., San Diego, CA, USA), and destained O/N using demineralized water. Protein amounts were estimated by densitometry using the Fiji software^86^ using a dilution series of the *Thermoascus aurantiacus* LPMO TaAA9A with a known concentrations (unpublished results). For *E. coli* expression, the protein titers were normalized by the culture volumes.

### Western blot analysis

Enzyme samples were handled as described in “SDS-PAGE analysis”, although instead of staining the gel the proteins were transferred to a nitrocellulose membrane using an iBlot Dry Blotting System (Invitrogen, Thermo Fisher Scientific, Waltham, MA, USA) at 20V for 7 minutes. The membrane was blocked with 2 % skim milk at 4 °C for at least 24 hours, washed three times for 5 minutes each in TBS-T (20 mM Tris-HCl pH 7.6, 150 mM NaCl, 0.1 % (v/v) Tween-20), and the membrane was incubated with primary antibody (anti-his tag, 1:1000, Merck Millipore, Merck KGaA, Darmstadt, Germany) diluted in 2 % skim milk for 1 hour. The washing steps were repeated, and the membrane was incubated with the secondary antibody (anti-Mouse-HRP IgG, 1:10,000, Sigma-Aldrich, Saint Louis, MO, USA) diluted in TBS-T for 1 hour. The washing steps were repeated again, and the protein:antibody complexes were visualized using Amersham ECL Prime Western Blotting Detection Reagent (GE Healthcare, Chicago, IL, USA).

## Acknowledgements

We thank Leila Lo Leggio for her inputs on the production of LPMOs in *E. coli*, Kenneth Jensen for his inputs on the production of LPMOs in *B. subtilis*, and Radina Tokin for inputs on the AZCL-HEC assay. This work was supported by the Novo Nordisk Foundation (Grant no. NNF17SA0027704).

