## Supplemental data for "LyGo: A platform for rapid screening of lytic polysaccharide monooxygenase production"

### Supplementary Figure S1

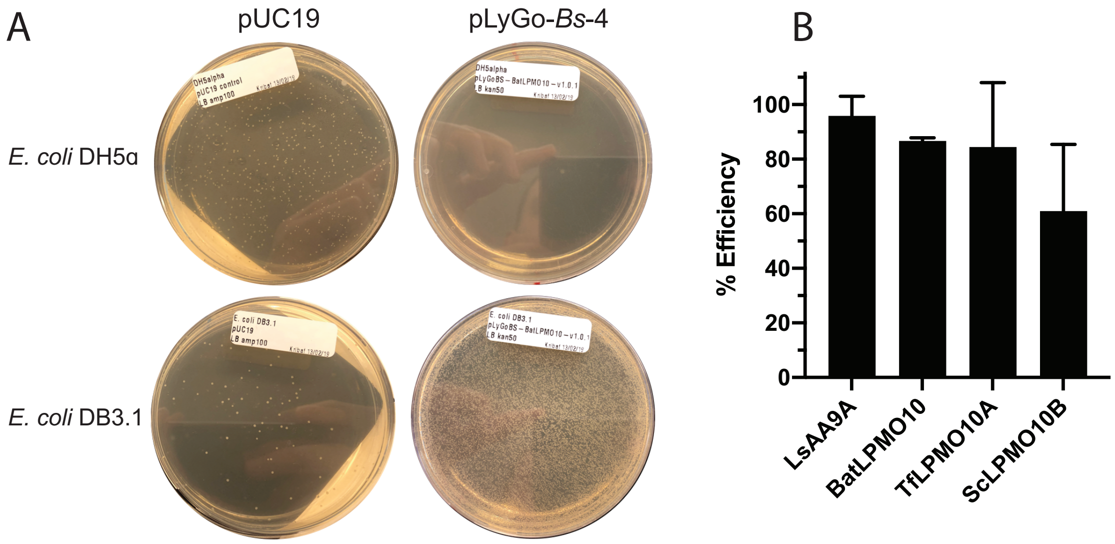


**Supplementary Figure S1. Counterselection and assembly efficiency of pLyGo vectors. (A)** Representative counterselection behavior of pLyGo vectors. pUC19 and pLyGo-*Bs*-4 were both transformed in *E. coli* DH5α and *E. coli* DB3.1 and selected on appropriate antibiotics. **(B)** Representative assembly efficiencies of different LyGo fragments and pLyGo-*Ec* vectors. The assembly efficiencies for the different LyGo fragments were estimated by LyGo cloning into pLyGo-*Ec*-2 through pLyGo-*Ec*-6. 8 resulting colonies from each reaction were tested by colony PCR, and the individual assembly efficiencies were calculated as the number of colonies exhibiting the correct PCR fragment divided by the number of colonies tested. The error bats indicate the standard deviation of biological triplicates.

### Supplementary Figure S2

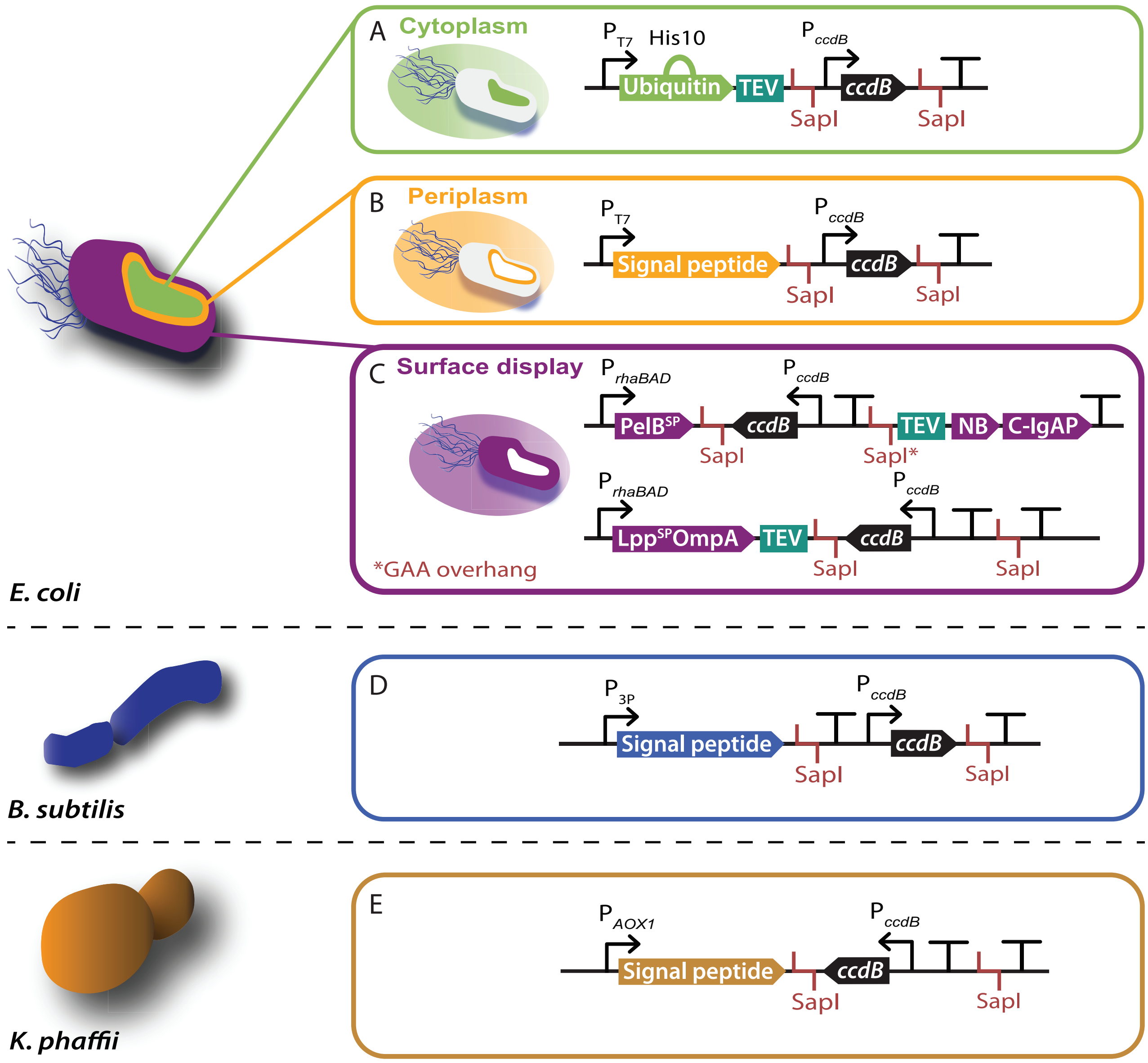


**Supplementary Figure S2. Architecture around the LyGo cassette for the different organisms.** Schematic overview of the area surrounding the LyGo cassette of the **(A)** cytoplasmic expression vectors for *E. coli*, **(B)** periplasmic expression vectors for *E. coli*, **(C)** surface display expression vectors for *E. coli* (top: C-terminal construct, bottom: N-terminal construct), **(D)** expression vectors for *B. subtilis*, **(E)** expression vectors for *K. phaffii*. Due to cloning issues, the design of the LyGo cassette was made in different versions. In all cases the expression promoter is situated upstream (to the left) of the LyGo cassette, with the reading direction from left to right.

### Supplementary Figure S3

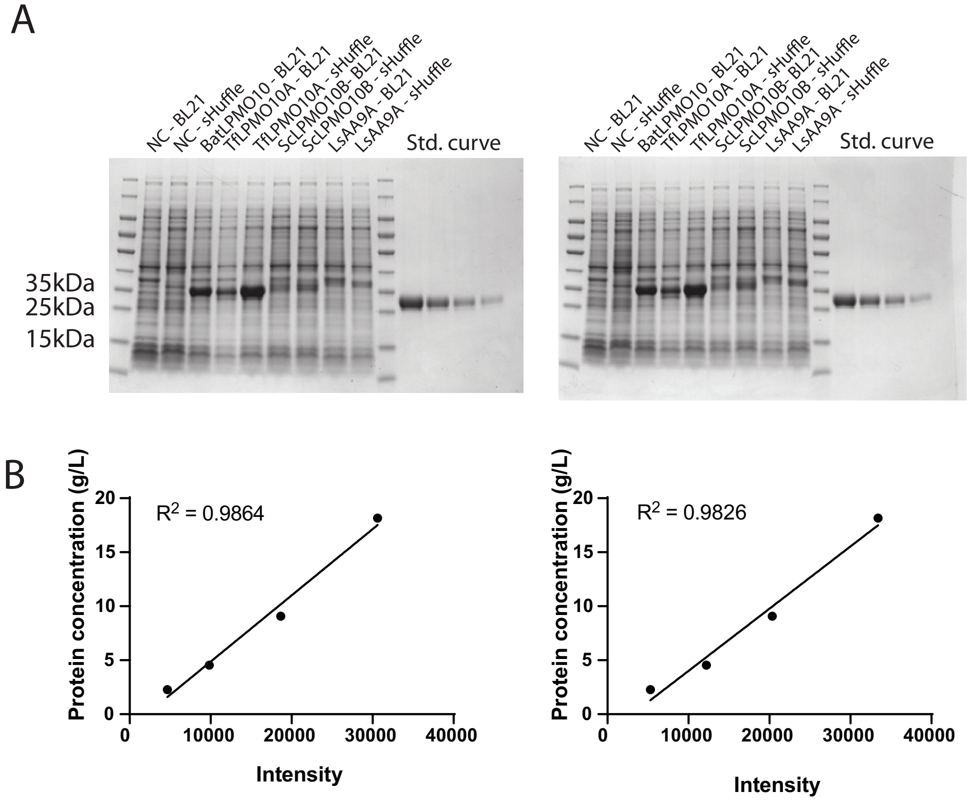


**Supplementary Figure S3. Quantification of protein concentration of samples expressed in the cytoplasm of *E. coli* BL21(DE3) and SHuffle strain.** **(A)** SDS-PAGE results for samples used for densitometry analysis of cytoplasmically expressed LPMOs. **(B)** Linear fit of the protein concentration as a function of pixel intensity, based on the standard curves. Pixel intensities were calculated in the Fiji software[1]. The concentration of the target protein constructs was calculated according to the linear regression fit and normalized per volume of culture collected.

### Supplementary Figure S4

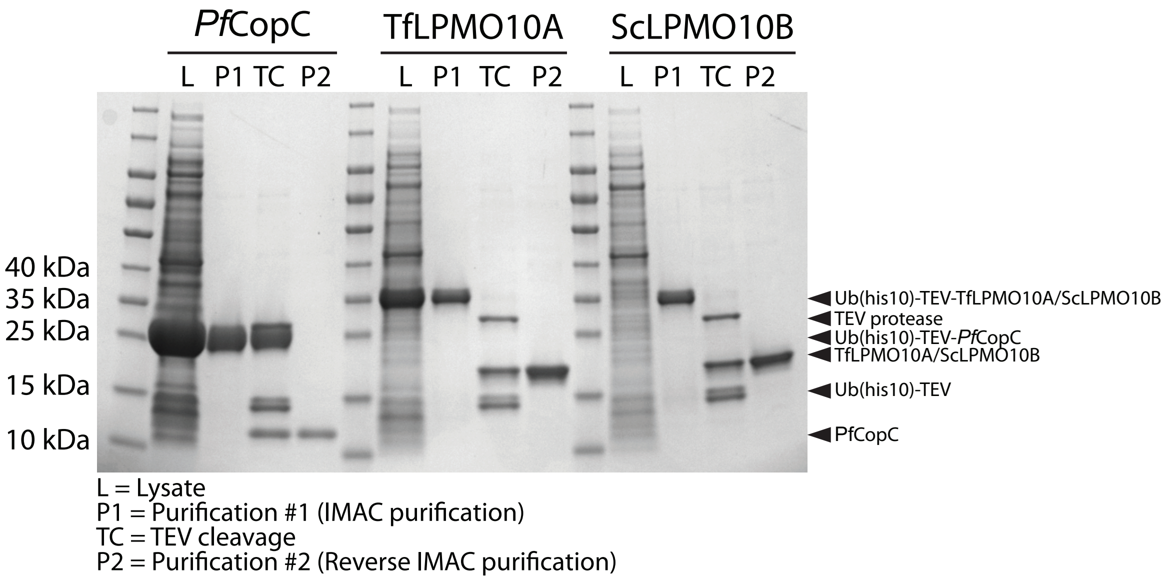


**Supplementary Figure S4. Production and purification of cytoplasmically produced proteins.** SDS-PAGE results for production and purification of PfCopC, TfLPMO10A, and ScLPMO10B using the Ub(his10)-tag (pLyGo-*Ec*-1).

### Supplementary Figure S5

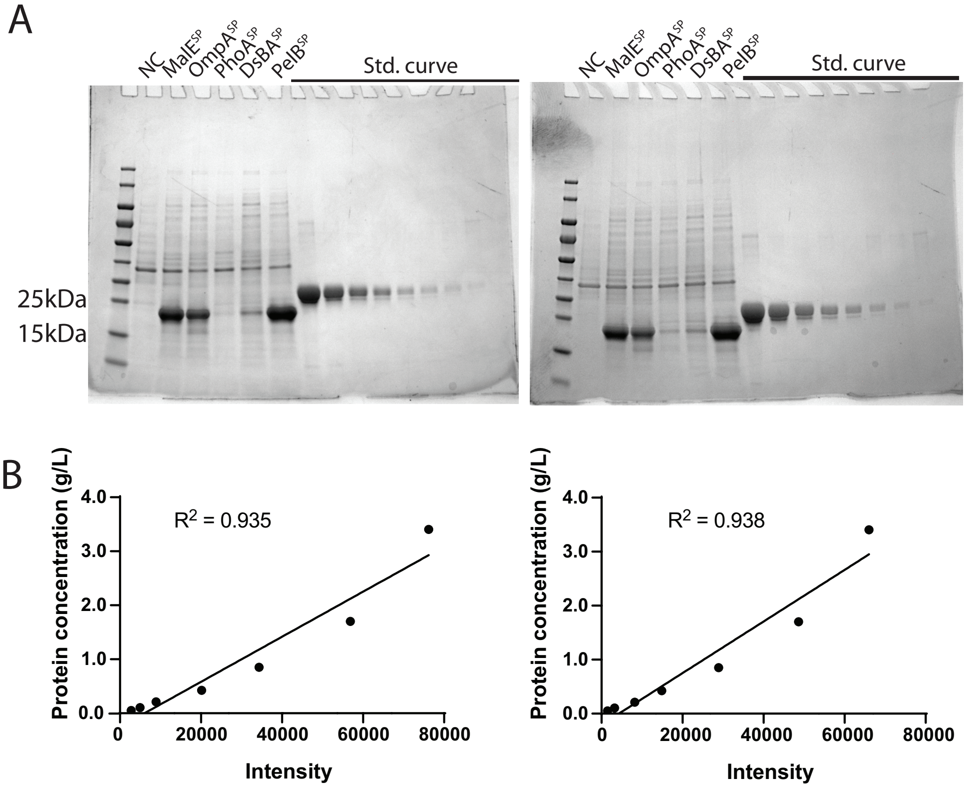


**Supplementary Figure S5. Quantification of protein concentration of BatLPMO10 samples expressed in the periplasm of *E. coli.*** **(A)** SDS-PAGE results for samples used for densitometry analysis of BatLPMO10 samples expressed in the periplasm of *E. coli.* **(B)** Linear fit of the protein concentration as a function of pixel intensity, based on the standard curves. Pixel intensities were calculated in the Fiji software[1]. The concentration of the target protein was calculated according to the linear regression fit and normalized per volume of culture collected.

### Supplementary Figure S6

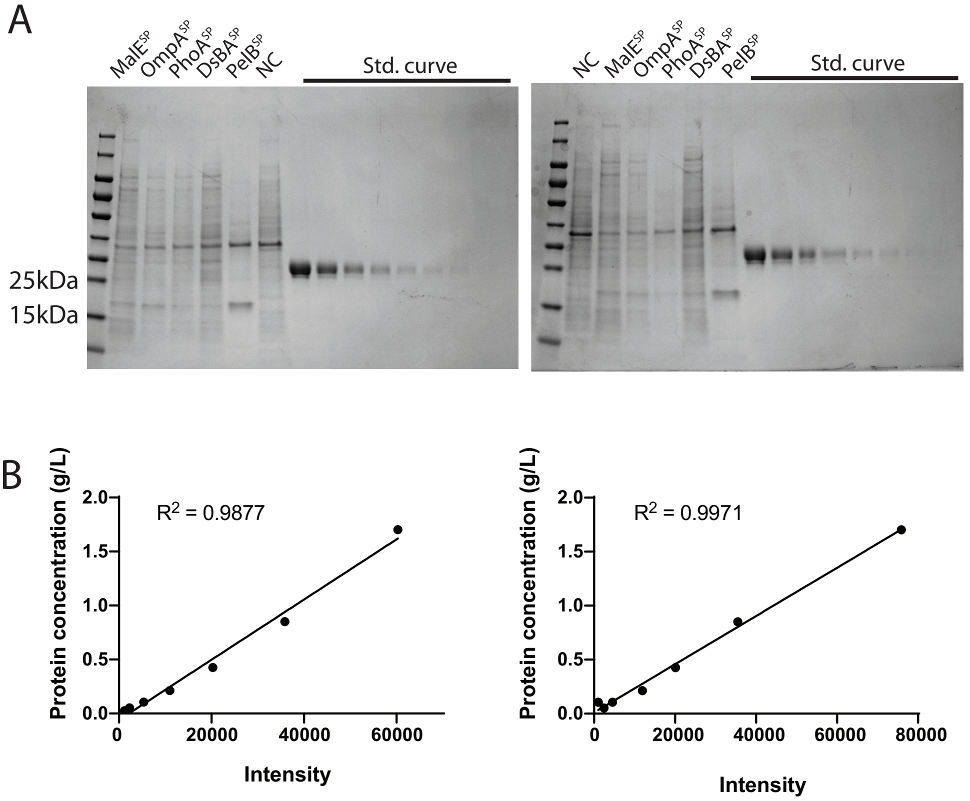


**Supplementary Figure S6. Quantification of protein concentration of TfLPMO10A samples expressed in the periplasm of *E. coli.*** **(A)** SDS-PAGE results for samples used for densitometry analysis of TfLPMO10A samples expressed in the periplasm of *E. coli.* **(B)** Linear fit of the protein concentration as a function of pixel intensity, based on the standard curves. Pixel intensities were calculated in the Fiji software[1]. The concentration of the target protein was calculated according to the linear regression fit and normalized per volume of culture collected.

### Supplementary Figure S7

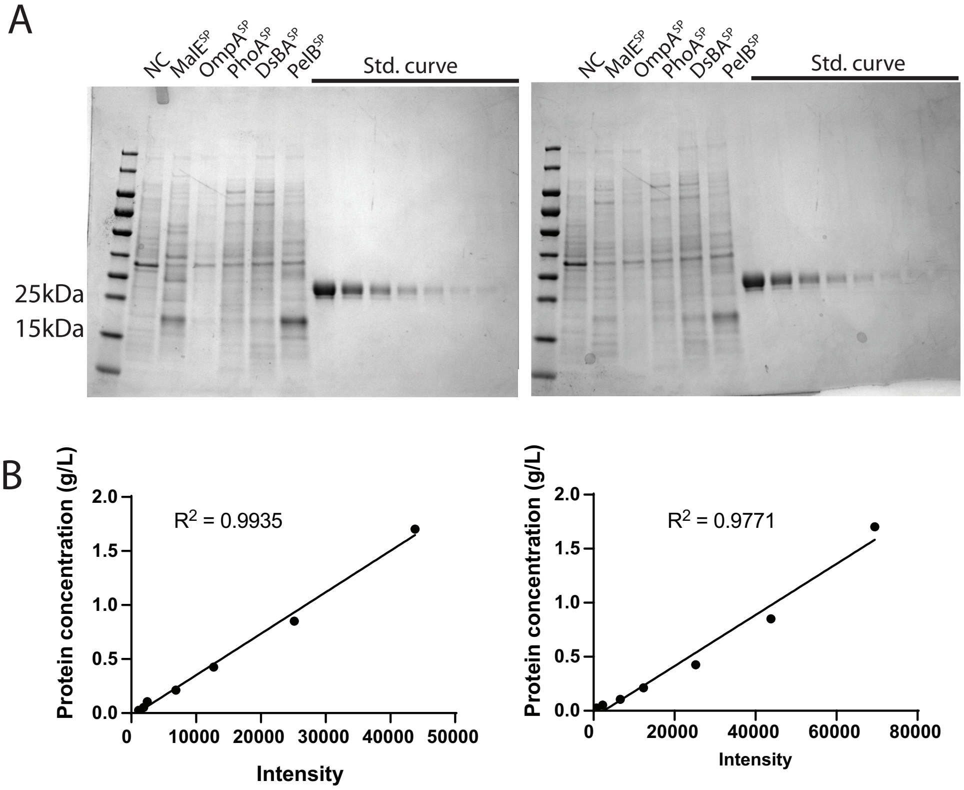


**Supplementary Figure S7. Quantification of protein concentration of ScLPMO10B samples expressed in the periplasm of *E. coli.* (A)** SDS-PAGE results for samples used for densitometry analysis of ScLPMO10B samples expressed in the periplasm of *E. coli.* **(B)** Linear fit of the protein concentration as a function of pixel intensity, based on the standard curves. Pixel intensities were calculated in the Fiji software[1]. The concentration of the target protein was calculated according to the linear regression fit and normalized per volume of culture collected.

### Supplementary Figure S8

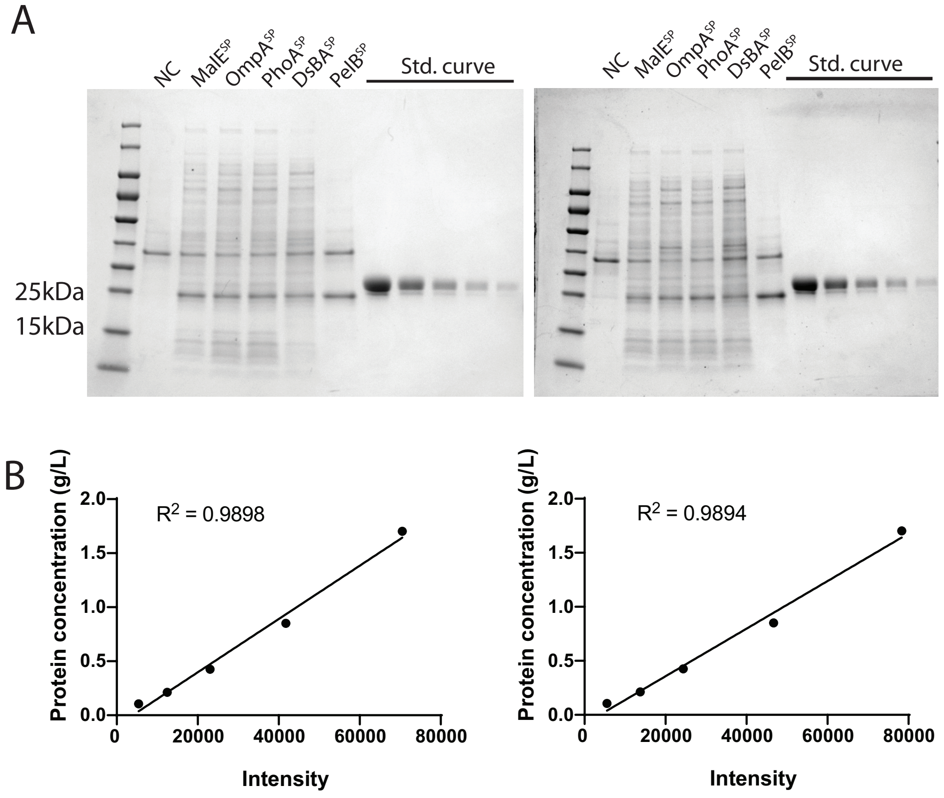


**Supplementary Figure S8. Quantification of protein concentration of LsAA9A samples expressed in the periplasm of *E. coli.*** **(A)** SDS-PAGE results for samples used for densitometry analysis of LsAA9A samples expressed in the periplasm of *E. coli.* **(B)** Linear fit of the protein concentration as a function of pixel intensity, based on the standard curves. Pixel intensities were calculated in the Fiji software[1]. The concentration of the target protein was calculated according to the linear regression fit and normalized per volume of culture collected.

### Supplementary Figure S9

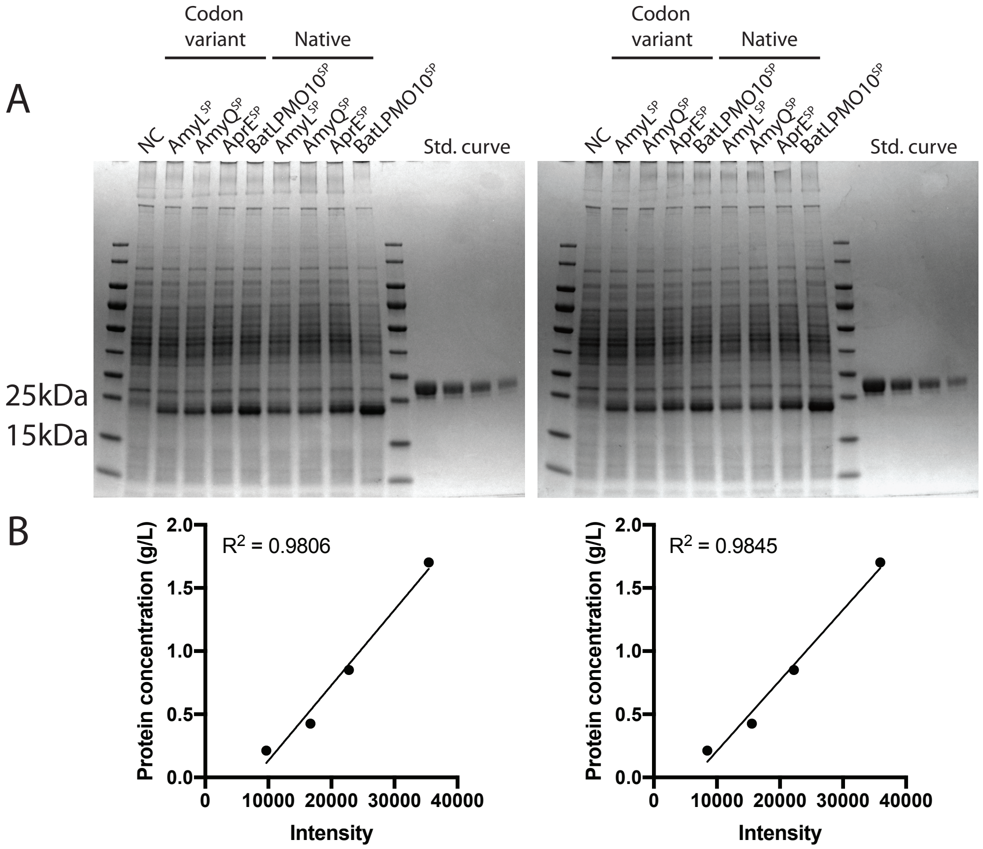


**Supplementary Figure S9. Quantification of protein concentration of BatLPMO10 in samples from *B. subtilis*. (A)** SDS-PAGE results for samples used for densitometry analysis of BatLPMO10 samples expressed in *B. subtilis.* **(B)** Linear fit of the protein concentration as a function of pixel intensity, based on the standard curves. Pixel intensities were calculated in the Fiji software[1]. The concentration of the target protein was calculated according to the linear regression fit.

### Supplementary Figure S10

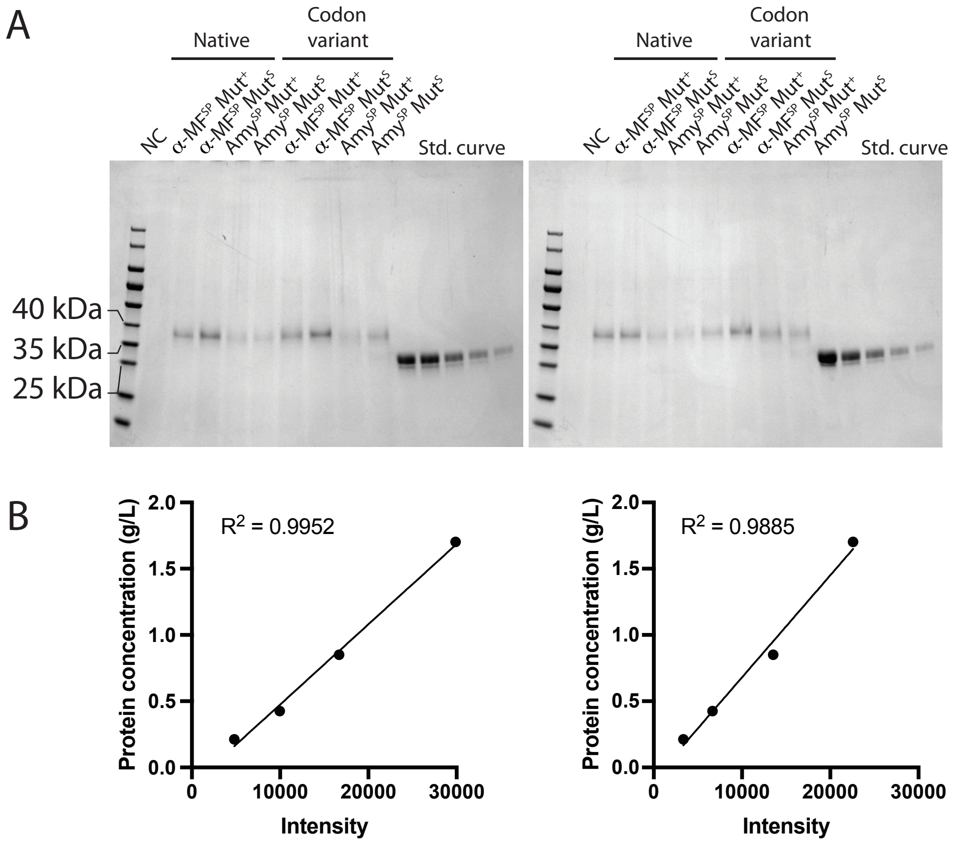


**Supplementary Figure S10. Quantification of protein concentration of LsAA9A in samples from *K. phaffii.*** **(A)** SDS-PAGE results for samples used for densitometry analysis of LsAA9A samples expressed in *K. phaffii.* **(B)** Linear fit of the protein concentration as a function of pixel intensity, based on the standard curves. Pixel intensities were calculated in the Fiji software[1]. The concentration of the target protein was calculated according to the linear regression fit.

### Supplementary Figure S11

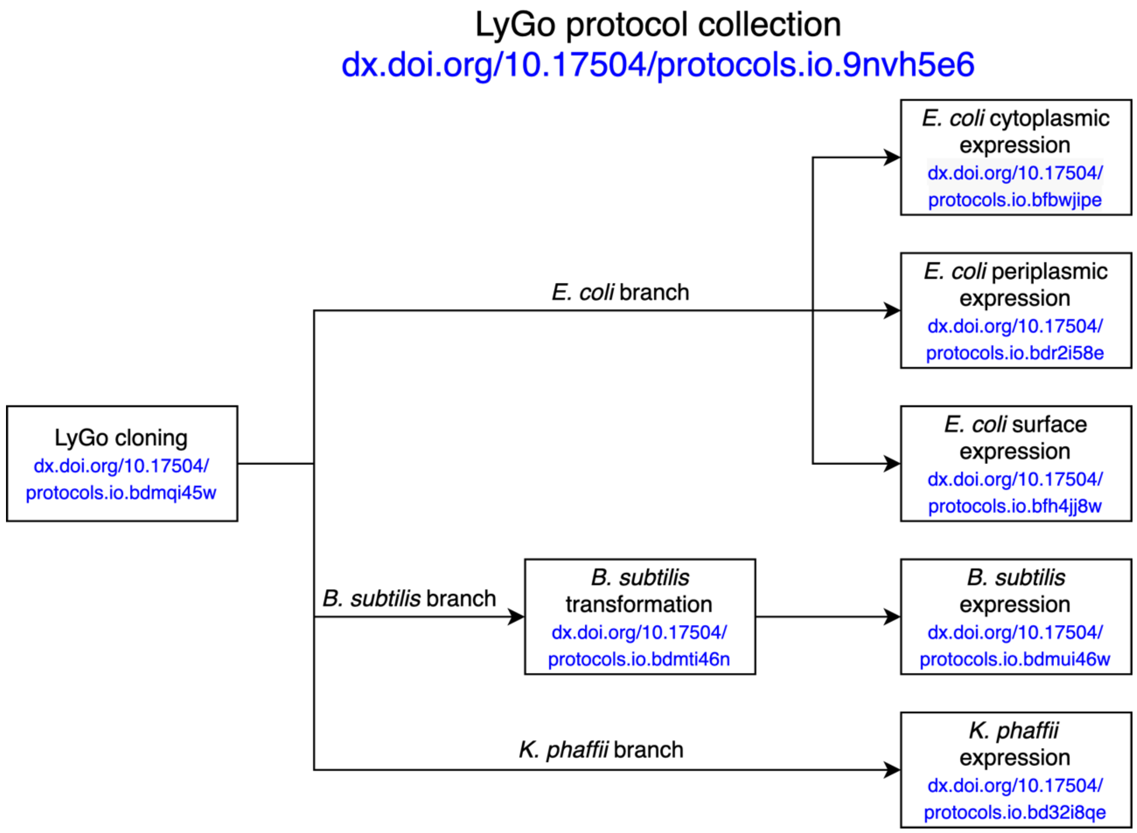


**Supplementary Figure S11. Workflow of protocols for cloning and expressing LPMOs.** Schematic overview of relationship between the provided protocols for cloning and expressing LPMOs in *E. coli*, *B. subtilis*, and *K. phaffii* using the LyGo platform. The DOI numbers provides direct links to the protocols.

### Supplementary Figure S12

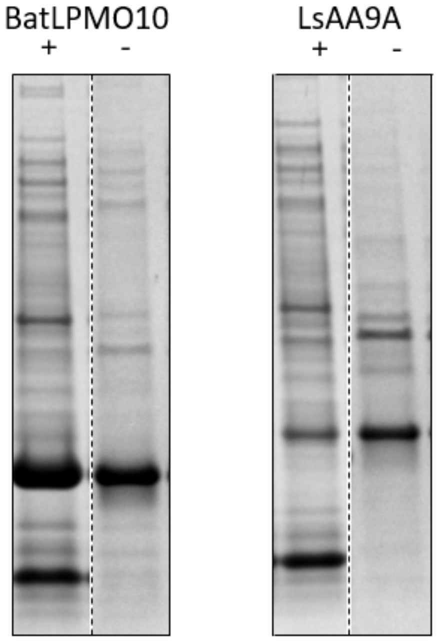


**Supplementary Figure S12. Comparison of different periplasmic extraction protocols.** Periplasmic extraction fractions of BatLPMO10 (left) and LsAA9A (right) using lysozyme (+) and without lysozyme (-) in TSE buffer.

### Supplementary Table S1

#### Supplementary Table S1. Strains used in this study

| Strain | Genotype | Source/Reference |
|---|---|---|
| *E. coli* DB3.1 | F$^{-}$ *gyrA*462 *endA1 glnV44* Δ(sr1-recA) *mcrB mrr hsdS20*(rB-, mB-) *ara14 galK2 lacY1 proA2 rpsL20*(SmR) *xyl5* Δ*leu mtl1* | *a* |
| *E. coli* NEB5α | *fhuA2* Δ(*argF-lacZ*)U169 *phoA glnV44* Φ80 Δ(*lacZ*)M15 *gyrA96 recA1 relA1 endA1 thi-1 hsdR17* | *b* |
| *E. coli* BL21(DE3) | F$^{-}$ *ompT gal dcm lon* $hsdS_B(r_B^- m_B^-)$ λ(DE3 [*lacI lacUV5-T7p07 ind1 sam7 nin5*]) [$malB^+$]$_{K-12}$(λ$^S$) | *c* |
| *E. coli* SHuffle | F′ *lac*, *pro*, *lacI*$^q$ / *Δ(ara-leu)7697 araD139 fhuA2 lacZ::T7 gene1 Δ(phoA)PvuII phoR ahpC* galE (or U) galK λatt*::pNEB3-r1-*cDsbC* (Spec$^R$, *lacI*$^q$) *ΔtrxB rpsL150*(Str$^R$) *Δgor Δ(malF)3* | Lab stock/[2] |
| *B. subtilis* KO7-S | Δ*nprE* Δ*aprE* Δ*epr* Δ*mpr* Δ*nprB* Δ*vpr* Δ*bpr* Δ*sigF* | BGSID1S145/*d* |
| *B. subtilis* KO7-S AmyL$^{SP}$-LsAA9A*nat* | Δ*nprE* Δ*aprE* Δ*epr* Δ*mpr* Δ*nprB* Δ*vpr* Δ*bpr* Δ*sigF* Δ*amyE*::P$_{3P}$-AmyL$^{SP}$-LsAA9A*nat*-His6 Cm$^R$ | This study |
| *B. subtilis* KO7-S AmyQ$^{SP}$-LsAA9A*nat* | Δ*nprE* Δ*aprE* Δ*epr* Δ*mpr* Δ*nprB* Δ*vpr* Δ*bpr* Δ*sigF* Δ*amyE*::P$_{3P}$-AmyQ$^{SP}$-LsAA9A*nat*-His6 Cm$^R$ | This study |
| *B. subtilis* KO7-S AprE$^{SP}$-LsAA9A*nat* | Δ*nprE* Δ*aprE* Δ*epr* Δ*mpr* Δ*nprB* Δ*vpr* Δ*bpr* Δ*sigF* Δ*amyE*::P$_{3P}$-AprE$^{SP}$-LsAA9A*nat*-His6 Cm$^R$ | This study |
| *B. subtilis* KO7-S BatLPMO10$^{SP}$-LsAA9A*nat* | Δ*nprE* Δ*aprE* Δ*epr* Δ*mpr* Δ*nprB* Δ*vpr* Δ*bpr* Δ*sigF* Δ*amyE*::P$_{3P}$-BatLPMO10$^{SP}$-LsAA9A*nat*-His6 Cm$^R$ | This study |
| *B. subtilis* KO7-S AmyL$^{SP}$-LsAA9A*opt* | Δ*nprE* Δ*aprE* Δ*epr* Δ*mpr* Δ*nprB* Δ*vpr* Δ*bpr* Δ*sigF* Δ*amyE*::P$_{3P}$-AmyL$^{SP}$-LsAA9A*opt*-His6 Cm$^R$ | This study |
| *B. subtilis* KO7-S AmyQ$^{SP}$-LsAA9A*opt* | Δ*nprE* Δ*aprE* Δ*epr* Δ*mpr* Δ*nprB* Δ*vpr* Δ*bpr* Δ*sigF* Δ*amyE*::P$_{3P}$-AmyQ$^{SP}$-LsAA9A*opt*-His6 Cm$^R$ | This study |
| *B. subtilis* KO7-S AprE$^{SP}$-LsAA9A*opt* | Δ*nprE* Δ*aprE* Δ*epr* Δ*mpr* Δ*nprB* Δ*vpr* Δ*bpr* Δ*sigF* Δ*amyE*::P$_{3P}$-AprE$^{SP}$-LsAA9A*opt*-His6 Cm$^R$ | This study |
| *B. subtilis* KO7-S BatLPMO10$^{SP}$-LsAA9A*opt* | Δ*nprE* Δ*aprE* Δ*epr* Δ*mpr* Δ*nprB* Δ*vpr* Δ*bpr* Δ*sigF* Δ*amyE*::P$_{3P}$-BatLPMO10$^{SP}$-LsAA9A*opt*-His6 Cm$^R$ | This study |
| *B. subtilis* KO7-S AmyL$^{SP}$-BatLPMO10*nat* | Δ*nprE* Δ*aprE* Δ*epr* Δ*mpr* Δ*nprB* Δ*vpr* Δ*bpr* Δ*sigF* Δ*amyE*::P$_{3P}$-AmyL$^{SP}$- BatLPMO10*nat* Cm$^R$ | This study |
| *B. subtilis* KO7-S AmyQ$^{SP}$- BatLPMO10*nat* | Δ*nprE* Δ*aprE* Δ*epr* Δ*mpr* Δ*nprB* Δ*vpr* Δ*bpr* Δ*sigF* Δ*amyE*::P$_{3P}$-AmyQ$^{SP}$-BatLPMO10*nat* Cm$^R$ | This study |
| *B. subtilis* KO7-S AprE$^{SP}$-BatLPMO10*nat* | Δ*nprE* Δ*aprE* Δ*epr* Δ*mpr* Δ*nprB* Δ*vpr* Δ*bpr* Δ*sigF* Δ*amyE*::P$_{3P}$-AprE$^{SP}$- BatLPMO10*nat* Cm$^R$ | This study |
| *B. subtilis* KO7-S BatLPMO10$^{SP}$-BatLPMO10*nat* | Δ*nprE* Δ*aprE* Δ*epr* Δ*mpr* Δ*nprB* Δ*vpr* Δ*bpr* Δ*sigF* Δ*amyE*::P$_{3P}$-BatLPMO10$^{SP}$-BatLPMO10*nat* Cm$^R$ | This study |
| *B. subtilis* KO7-S AmyL$^{SP}$- BatLPMO10*opt* | Δ*nprE* Δ*aprE* Δ*epr* Δ*mpr* Δ*nprB* Δ*vpr* Δ*bpr* Δ*sigF* Δ*amyE*::P$_{3P}$-AmyL$^{SP}$- BatLPMO10*opt* Cm$^R$ | This study |
| *B. subtilis* KO7-S AmyQ$^{SP}$- BatLPMO10*opt* | Δ*nprE* Δ*aprE* Δ*epr* Δ*mpr* Δ*nprB* Δ*vpr* Δ*bpr* Δ*sigF* Δ*amyE*::P$_{3P}$-AmyQ$^{SP}$-BatLPMO10*opt* Cm$^R$ | This study |
| *B. subtilis* KO7-S AprE$^{SP}$-BatLPMO10*opt* | Δ*nprE* Δ*aprE* Δ*epr* Δ*mpr* Δ*nprB* Δ*vpr* Δ*bpr* Δ*amyE*::P$_{3P}$-AprE$^{SP}$-BatLPMO10*opt* Cm$^R$ | This study |

| | | |
|---|---|---|
| *B. subtilis* KO7-S BatLPMO10$^{SP}$-BatLPMO10*opt* | Δ*nprE* Δ*aprE* Δ*epr* Δ*mpr* Δ*nprB* Δ*vpr* Δ*bpr* *amyE*::$P_{3P}$-BatLPMO10$^{SP}$-BatLPMO10*opt* Cm$^{R}$ | This study |
| *K. phaffii (P. pastoris)* GS115 | *his4* | *a* |
| *K. phaffii* GS115 α-MF$^{SP}$-LsAA9A*nat* Mut$^{S}$ | *aox1*::$P_{AOX1}$-α-MF$^{SP}$-LsAA9A*nat*-His6-*HIS4* | This study |
| *K. phaffii* GS115 α-MF$^{SP}$-LsAA9A*nat* Mut$^{+}$ | *his4*::$P_{AOX1}$-α-MF$^{SP}$-LsAA9A*nat*-His6-*HIS4* | This study |
| *K. phaffii* GS115 Amy$^{SP}$-LsAA9A*nat* Mut$^{S}$ | *aox1*::$P_{AOX1}$-Amy$^{SP}$-LsAA9A*na*-His6*t*-*HIS4* | This study |
| *K. phaffii* GS115 Amy$^{SP}$-LsAA9A*nat* Mut$^{+}$ | *his4*::$P_{AOX1}$-Amy$^{SP}$-LsAA9A*nat*-His6-*HIS4* | This study |
| *K. phaffii* GS115 α-MF$^{SP}$-LsAA9A*opt* Mut$^{S}$ | *aox1*::$P_{AOX1}$-α-MF$^{SP}$-LsAA9A*op*-His6*t*-*HIS4* | This study |
| *K. phaffii* GS115 α-MF$^{SP}$-LsAA9A*opt* Mut$^{+}$ | *his4*::$P_{AOX1}$-α-MF$^{SP}$-LsAA9A*opt*-His6-*HIS4* | This study |
| *K. phaffii* GS115 Amy$^{SP}$-LsAA9A*opt* Mut$^{S}$ | *aox1*::$P_{AOX1}$-Amy$^{SP}$-LsAA9A*opt*-His6-*HIS4* | This study |
| *K. phaffii* GS115 Amy$^{SP}$-LsAA9A*opt* Mut$^{+}$ | *his4*::$P_{AOX1}$-Amy$^{SP}$-LsAA9A*opt*-His6-*HIS4* | This study |

[a]Thermo Fisher Scientific, Waltham, MA, USA; [b]NEB, Ipswich, MA, USA; [c]Novagen, Merck KGaA, Darmstadt, Germany; [d]Bacillus Genetic Stock Center, Columbus, OH, USA

### Supplementary Table S2

**Supplementary Table S2. Plasmids used in this study**

| Plasmid | Description | Source/Reference |
|---|---|---|
| pET28a(+)-MalE$^{SP}$-*bla* | vector encoding *malE* signal sequence and *bla* gene, Km$^{R}$ | [3] |
| pET28a(+)-OmpA$^{SP}$-*bla* | vector encoding *ompA* signal sequence and *bla* gene, Km$^{R}$ | [3] |
| pET28a(+)-PhoA$^{SP}$-*bla* | vector encoding *phoA* signal sequence and *bla* gene, Km$^{R}$ | [3] |
| pET28a(+)-DsbA$^{SP}$-*bla* | vector encoding *dsbA* signal sequence and *bla* gene, Km$^{R}$ | [3] |
| pET28a(+)-PelB$^{SP}$-*bla* | vector encoding *pelB* signal sequence and *bla* gene, Km$^{R}$ | [3] |
| pMSB881 | vector encoding super folder GFP, Amp$^{R}$ | Lab stock |
| pTEVprotease | vector encoding the His-tagged TEV protease, Amp$^{R}$ | Provided by Lars Ellgaard |
| pET39-Ub(His10)-TEV-G101 | vector encoding Ubiquitin with an internal His10-tag, Km$^{R}$ | Provided by Lars Ellgaard |
| pBAD42-Lpp$^{SP}$-OmpA-NB | vector encoding Lpp-OmpA-NB surface expression construct, Km$^{R}$ | [4] |

| | | |
|---|---|---|
| pBAD42-NB-C-IgAP | vector encoding NB-C-IgAP surface expression construct, $\mathrm{Km}^{R}$ | [4] |
| pBSc243C | Bacillus SEVA sibling vector carrying *lacZ*α in the SEVA cargo site, $\mathrm{Ap}^{R}$ (*E.* coli), $\mathrm{Cm}^{R}$ (*B. subtilis*) | [5] |
| pACYC-SL3m-*ccdB* | vector encoding *ccdB* gene, $\mathrm{Cm}^{R}$ | Lab stock |
| pBAD42-$\mathrm{Lpp}^{SP}$-OmpA-TEV-NB | vector encoding *Lpp-OmpA* surface expression anchor and nanobody, $\mathrm{Km}^{R}$ | This study |
| pLyGo-*Bs*-NC | pLyGo-*Bs* vector where the CDS is a start codon and a stop codon, used as negative control, | This study |

| LyGo nomenclature | Classical nomenclature | Description | Source/Reference |
|---|---|---|---|
| ***E. coli* expression vectors** | | | |
| pLyGo-*Ec*-1-BatLPMO10 | pET39b-Ub(His10)-TEV*-BatLPMO10 | vector containing a ubiquitin solubility tag and BatLPMO10 | This study |
| pLyGo-*Ec*-1-TfLPMO10A | pET39b-Ub(His10)-TEV*-TfLPMO10A | vector containing a ubiquitin solubility tag and TfLPMO10A | This study |
| pLyGo-*Ec*-1-ScLPMO10B | pET39b-Ub(His10)-TEV*-ScLPMO10B | vector containing a ubiquitin solubility tag and LsAA9A | This study |
| pLyGo-*Ec*-1-LsAA9A | pET39b-Ub(His10)-TEV*-LsAA9A | vector containing a ubiquitin solubility tag and LsAA9A | This study |
| pLyGo-*Ec*-1-PfCopC | pET39b-Ub(His10)-TEV*-PfCopC | vector containing a ubiquitin solubility tag and PfCopC | This study |
| pLyGo-*Ec*-2-BatLPMO10 | pET28a(+)-$\mathrm{MalE}^{SP}$-BatLPMO10 | vector encoding *malE* signal sequence and BatLPMO10, $\mathrm{Km}^{R}$ | This study |
| pLyGo-*Ec*-3-BatLPMO10 | pET28a(+)-$\mathrm{OmpA}^{SP}$-BatLPMO10 | vector encoding *ompA* signal sequence and BatLPMO10, $\mathrm{Km}^{R}$ | This study |
| pLyGo-*Ec*-4-BatLPMO10 | pET28a(+)-$\mathrm{PhoA}^{SP}$-BatLPMO10 | vector encoding *phoA* signal sequence and BatLPMO10, $\mathrm{Km}^{R}$ | This study |
| pLyGo-*Ec*-5-BatLPMO10 | pET28a(+)-$\mathrm{DsbA}^{SP}$-BatLPMO10 | vector encoding *dsbA* signal sequence and BatLPMO10, $\mathrm{Km}^{R}$ | This study |
| pLyGo-*Ec*-6-BatLPMO10 | pET28a(+)-$\mathrm{PelB}^{SP}$-BatLPMO10 | vector encoding *pelB* signal sequence and BatLPMO10, $\mathrm{Km}^{R}$ | This study |
| pLyGo-*Ec*-2-TfLPMO10A | pET28a(+)-$\mathrm{MalE}^{SP}$-TfLPMO10A | vector encoding *malE* signal sequence and TfLPMO10A, $\mathrm{Km}^{R}$ | This study |
| pLyGo-*Ec*-3-TfLPMO10A | pET28a(+)-$\mathrm{OmpA}^{SP}$-TfLPMO10A | vector encoding *ompA* signal sequence and TfLPMO10A, $\mathrm{Km}^{R}$ | This study |
| pLyGo-*Ec*-4-TfLPMO10A | pET28a(+)-$\mathrm{PhoA}^{SP}$-TfLPMO10A | vector encoding *phoA* signal sequence and TfLPMO10A, $\mathrm{Km}^{R}$ | This study |
| pLyGo-*Ec*-5-TfLPMO10A | pET28a(+)-$\mathrm{DsbA}^{SP}$-TfLPMO10A | vector encoding *dsbA* signal sequence and TfLPMO10A, $\mathrm{Km}^{R}$ | This study |
| pLyGo-*Ec*-6-TfLPMO10A | pET28a(+)-$\mathrm{PelB}^{SP}$-TfLPMO10A | vector encoding *pelB* signal sequence and TfLPMO10A, $\mathrm{Km}^{R}$ | This study |
| pLyGo-*Ec*-2-ScLPMO10B | pET28a(+)-$\mathrm{MalE}^{SP}$-ScLPMO10B | vector encoding *malE* signal sequence and ScLPMO10B, $\mathrm{Km}^{R}$ | This study |
| pLyGo-*Ec*-3-ScLPMO10B | pET28a(+)-$\mathrm{OmpA}^{SP}$-ScLPMO10B | vector encoding *ompA* signal sequence and ScLPMO10B, $\mathrm{Km}^{R}$ | This study |

| | | | |
|---|---|---|---|
| pLyGo-*Ec*-4-ScLPMO10B | pET28a(+)-PhoA$^{SP}$-ScLPMO10B | vector encoding *phoA* signal sequence and ScLPMO10B, Km$^{R}$ | This study |
| pLyGo-*Ec*-5-ScLPMO10B | pET28a(+)-DsbA$^{SP}$-ScLPMO10B | vector encoding *dsbA* signal sequence and ScLPMO10B, Km$^{R}$ | This study |
| pLyGo-*Ec*-6-ScLPMO10B | pET28a(+)-PelB$^{SP}$-ScLPMO10B | vector encoding *pelB* signal sequence and ScLPMO10B, Km$^{R}$ | This study |
| pLyGo-*Ec*-2-LsAA9A | pET28a(+)-MalE$^{SP}$-LsAA9A | vector encoding *malE* signal sequence and LsAA9A, Km$^{R}$ | This study |
| pLyGo-*Ec*-3-LsAA9A | pET28a(+)-OmpA$^{SP}$-LsAA9A | vector encoding *ompA* signal sequence and LsAA9A, Km$^{R}$ | This study |
| pLyGo-*Ec*-4-LsAA9A | pET28a(+)-PhoA$^{SP}$-LsAA9A | vector encoding *phoA* signal sequence and LsAA9A, Km$^{R}$ | This study |
| pLyGo-*Ec*-5-LsAA9A | pET28a(+)-DsbA$^{SP}$-LsAA9A | vector encoding *dsbA* signal sequence and LsAA9A, Km$^{R}$ | This study |
| pLyGo-*Ec*-6-LsAA9A | pET28a(+)-PelB$^{SP}$-LsAA9A | vector encoding *pelB* signal sequence and LsAA9A, Km$^{R}$ | This study |
| pLyGo-*Ec*-7-LsAA9A | pBAD42-LsAA9A-his8-TEV-NB-C-IgAP | vector encoding a single domain antibody (nanobody, NB), C-IgAP surface expression anchor and LsAA9A, Km$^{R}$ | This study |
| pLyGo-*Ec*-8-LsAA9A-His8 | pBAD42-Lpp$^{SP}$-OmpA-TEV*-LsAA9A-His8 | vector encoding Lpp-OmpA surface expression anchor and LsAA9A, Km$^{R}$ | This study |
| pLyGo-*Ec*-8-ScLPMO10-His8 | pBAD42-Lpp$^{SP}$-OmpA-TEV*-ScLPMO10B-His8 | vector encoding Lpp-OmpA surface expression anchor and ScLPMO10B, Km$^{R}$ | This study |
| ***B. subtilis* expression vectors** | | | |
| pLyGo-*Bs*-1-LsAA9A*nat*-His6 | pBS293C-amyE-$P_{3P}$-AmyL$^{SP}$-LsAA9A*nat*-His6 | Vector encoding the *amyL* signal peptide and LsAA9A*nat*, Km$^{R}$ (*E. coli*), Cm$^{R}$ (*B. subtilis*) | This study |
| pLyGo-*Bs*-2-LsAA9A*nat*-His6 | pBS293C-amyE-$P_{3P}$-AmyQ$^{SP}$-LsAA9A*nat*-His6 | Vector encoding the *amyQ* signal peptide and LsAA9A*nat*, Km$^{R}$ (*E. coli*), Cm$^{R}$ (*B. subtilis*) | This study |
| pLyGo-*Bs*-3-LsAA9A*nat*-His6 | pBS293C-amyE-$P_{3P}$-AprE$^{SP}$-LsAA9A*nat*-His6 | Vector encoding the *aprE* signal peptide and LsAA9A*nat*, Km$^{R}$ (*E. coli*), Cm$^{R}$ (*B. subtilis*) | This study |
| pLyGo-*Bs*-4-LsAA9A*nat*-His6 | pBS293C-amyE-$P_{3P}$-BatLPMO10$^{SP}$-LsAA9A*nat*-His6 | Vector encoding the BatLPMO10 signal peptide and LsAA9A*nat*, Km$^{R}$ (*E. coli*), Cm$^{R}$ (*B. subtilis*) | This study |
| pLyGo-*Bs*-1-LsAA9A*opt*-His6 | pBS293C-amyE-$P_{3P}$-AmyL$^{SP}$-LsAA9A*opt*-His6 | Vector encoding the *amyL* signal peptide and LsAA9A*opt*, Km$^{R}$ (*E. coli*), Cm$^{R}$ (*B. subtilis*) | This study |
| pLyGo-*Bs*-2-LsAA9A*opt*-His6 | pBS293C-amyE-$P_{3P}$-AmyQ$^{SP}$-LsAA9A*opt*-His6 | Vector encoding the *amyQ* signal peptide and LsAA9A*opt*, Km$^{R}$ (*E. coli*), Cm$^{R}$ (*B. subtilis*) | This study |
| pLyGo-*Bs*-3-LsAA9A*opt*-His6 | pBS293C-amyE-$P_{3P}$-AprE$^{SP}$-LsAA9A*opt*-His6 | Vector encoding the *aprE* signal peptide and LsAA9A*opt*, Km$^{R}$ (*E. coli*), Cm$^{R}$ (*B. subtilis*) | This study |

| | | | |
|---|---|---|---|
| pLyGo-*Bs*-4-LsAA9A*opt*-His6 | pBS293C-amyE-$P_{3P}$-BatLPMO10$^{SP}$-LsAA9A*opt*-His6 | Vector encoding the BatLPMO10 signal peptide and LsAA9A*opt*, $Km^R$ (*E. coli*), $Cm^R$ (*B. subtilis*) | This study |
| pLyGo-*Bs*-1-BatLPMO10*nat*-His6 | pBS293C-amyE-$P_{3P}$-AmyL$^{SP}$-BatLPMO10*nat*-His6 | Vector encoding the *amyL* signal peptide and BatLPMO10*nat*, $Km^R$ (*E. coli*), $Cm^R$ (*B. subtilis*) | This study |
| pLyGo-*Bs*-2-BatLPMO10*nat* -His6 | pBS293C-amyE-$P_{3P}$-AmyQ$^{SP}$- BatLPMO10*nat*-His6 | Vector encoding the *amyQ* signal peptide and BatLPMO10*nat*, $Km^R$ (*E. coli*), $Cm^R$ (*B. subtilis*) | This study |
| pLyGo-*Bs*-3-BatLPMO10*nat* -His6 | pBS293C-amyE-$P_{3P}$-AprE$^{SP}$- BatLPMO10*nat*-His6 | Vector encoding the *aprE* signal peptide and BatLPMO10*nat*, $Km^R$ (*E. coli*), $Cm^R$ (*B. subtilis*) | This study |
| pLyGo-*Bs*-4-BatLPMO10*nat* -His6 | pBS293C-amyE-$P_{3P}$-BatLPMO10$^{SP}$-BatLPMO10*nat*-His6 | Vector encoding the BatLPMO10 signal peptide and BatLPMO10*nat*, $Km^R$ (*E. coli*), $Cm^R$ (*B. subtilis*) | This study |
| pLyGo-*Bs*-1-BatLPMO10*opt* | pBS293C-amyE-$P_{3P}$-AmyL$^{SP}$- BatLPMO10*opt* | Vector encoding the *amyL* signal peptide and BatLPMO10*opt*, $Km^R$ (*E. coli*), $Cm^R$ (*B. subtilis*) | This study |
| pLyGo-*Bs*-2-BatLPMO10*opt* | pBS293C-amyE-$P_{3P}$-AmyQ$^{SP}$- BatLPMO10*opt* | Vector encoding the *amyQ* signal peptide and BatLPMO10*opt*, $Km^R$ (*E. coli*), $Cm^R$ (*B. subtilis*) | This study |
| pLyGo-*Bs*-3-BatLPMO10*opt* | pBS293C-amyE-$P_{3P}$-AprE$^{SP}$- BatLPMO10*opt* | Vector encoding the *aprE* signal peptide and BatLPMO10*opt*, $Km^R$ (*E. coli*), $Cm^R$ (*B. subtilis*) | This study |
| pLyGo-*Bs*-4-BatLPMO10*opt* | pBS293C-amyE-$P_{3P}$-BatLPMO10$^{SP}$-BatLPMO10*opt* | Vector encoding the BatLPMO10 signal peptide and BatLPMO10*opt*, $Km^R$ (*E. coli*), $Cm^R$ (*B. subtilis*) | This study |
| ***K. phaffii* expression vectors** | | | |
| pLyGo-*Kp*-1-LsAA9A*nat*-His6 | pPIC9K-α-MF$^{SP}$-LsAA9A*nat*-His6 | vector encoding α-*MF* signal sequence and LsAA9A*nat,* $Amp^R$ (*E. coli*), $Km^R$, *HIS4* (*K. phaffii*) | This study |
| pLyGo-*Kp*-1-LsAA9A*opt*-His6 | pPIC9K-α-MF$^{SP}$-LsAA9A*opt*-His6 | vector encoding α-*MF* signal sequence and LsAA9A*opt,* $Amp^R$ (*E. coli*), $Km^R$, *HIS4* (*K. phaffii*) | This study |
| pLyGo-*Kp*-2-LsAA9A*nat*-His6 | pPIC9K-Amy$^{SP}$-LsAA9A*nat*-His6 | vector encoding α-amylase signal sequence and LsAA9A*nat,* $Amp^R$ (*E. coli*), $Km^R$, *HIS4* (*K. phaffii*) | This study |
| pLyGo-*Kp*-2-LsAA9A*opt*-His6 | pPIC9K-Amy$^{SP}$-LsAA9A*opt*-His6 | vector encoding α-amylase signal sequence and LsAA9A*opt,* $Amp^R$ (*E. coli*), $Kan^R$, *HIS4* (*K. phaffii*) | This study |

### Supplementary Table S3

**Supplementary Table S3. Oligonucleotides used in this study**

| No. | Name | Sequence (5' --> 3') |
|---|---|---|
| **Cloning oligonucleotides** | | |
| 1 | FW_universal_BB | ATCCGGCUGCTAACAAAGCCCG |

| | | |
|---|---|---|
| 2 | FW_ccdB | ATCGAAGAGCUACTAAAAGCCAGATAACAGTATGCGTATTTG |
| 3 | RV_ccdB_stop_NoGly | AGCCGGAUCTTACGAAGAGCCTATTCCCC |
| 4 | RW_DsBA | AGCTCTTCGAUGCGCCGATGCGCTAAACGC |
| 5 | RW_MalE | AGCTCTTCGAUGGGCGAGAGCCGAGGC |
| 6 | RW_OmpA | AGCTCTTCGAUGGGCCTGCGCTACGGTAGC |
| 7 | RW_PhoA | AGCTCTTCGAUGGGCTTTTGTCACAGGGGTAAACAGTAAC |
| 8 | RW_pelB_BB | AGCTCTTCGAUGGGCCATCGCCGGCTG |
| 9 | FW_BOM | ATGCGGTAUTTTCTCCTTACGCATCTG |
| 10 | RW_sapI_removal | ATACCGCAUCAGGCCCTCTTCCGCTTCCTCGCTCACTGAC |
| 11 | FW_TEV_GFP | AGGCTCTUCGGGTGGATCCGAAAACCTGTACTTCCAGGGTCAATTCAG |
| 12 | RW_TEV_GFPfr | AGCCGGAUCTCAGTGGTGGTGGTGGTGATGATGATG |
| 13 | RW_ccdB_nostop | AAGAGCCUATTCCCCAGAACATCAGGTTAATGGCG |
| 14 | -gfp+HIS F | ATCATCAUCACCACCACCACCACTG |
| 15 | Pfu-to-Waldo_vrev | ATTGACCCUGGAAGTACAGGTTTTC |
| 16 | FW_TEV+GFPsf | AGGGTCAAUTCATGCGTAAAGGCGAGGAGC |
| 17 | RW_GFPsf_HIS | ATGATGAUGGGCCGCAAGCTTTTTGTACAGTTCATCCATACCATGCGT |
| 18 | RV_newTEVoptTIR | AGCTCTTCGAUGTTGGAAGTACAGGTTTTCAGCCATACAACTATC |
| 19 | RV_USER_pET39_BB | ATCCGGCUCCACCGCTGAGCAATAACT |
| 20 | FW_USER_pET39_BB | AAGTACAGGTUTTCACCGCCAGAAGCGTCA |
| 21 | pBR322_USER_rev | ACCGCCAGUGCATGTTCATCATCAGTAACCC |
| 22 | pBR322_KanR_USER_fwd | AGAAGACTUAGGAACTATATCCGGATTGGC |
| 23 | amyE_up_USER_fwd | ACTGGCGGUATGTTTGCAAAACGATTCAA |
| 24 | amyE_up_USER_rev | AGACGAUCGATCAGACCAGTTTTTAATTTG |
| 25 | amyE_down_USER_fwd | ATTACGATUGCGCCTCATGTTTTCC |
| 26 | amyE_down_USER_rev | AAGTCTTCUTCAATGGGGAAGAGAACC |
| 27 | pBS_Cargo_AB_USER_fwd | ATCGTCUCAGCGAGGC |
| 28 | PromoterSwap_GibsonAssembly_rev | ATAGACGCATAAAGTGTAAAGCC |
| 29 | PromoterSwap_GibsonAssembly_fwd | GATTAACTAATAAGGAGGACAAACATG |
| 30 | pBS_USER_rev2 | ATAGACGCAUAAAGTGTAAAGCCTGG |
| 31 | SP_amyL_fwd | ATCTTCTTGCUGCCTCATTCTGCAGCAGCGGCGATGAGTAAAGGCGAAGAGCT |
| 32 | SP_amyL_rev2_USER | AGCAAGAAGAUGAGCGCAAATAACAGCGGCAGCAATCGGGCGTAAAGCCGTTTTTGTTGTTTCATGTTTGTCCTCCTTATTAGTTAATC |
| 33 | SP_amyQ_fwd | ACGCTGTUATTTGTCAGTTTGCCGATTACAAAAACATCAGCCATGAGTAAAGGCGAAGAGCT |
| 34 | SP_amyQ_rev2_USER | AACAGCGUGCACATAAGCACAAGTCTGAACGAAACTGTCCGCTTTCGTTTTTGAATCATGTTTGTCCTCCTTATTAGTTAATC |
| 35 | SP_aprE_fwd | ATTTCTGTUGCTTTTAGTTCATCGATCGCATCGGCTGCTATGAGTAAAGGCGAAGAGCT |
| 36 | SP_aprE_rev_USER | AACAGAAAUGAGTAGTGCGGTGCTTGCGACAATTTTCCCCAACGGTTTCTTCATGTTTGTCCTCCTTATTAGTTAATC |
| 37 | BatLPMO10_SP_USER_fwd | AGGCGTTGUTCTAACTGCTGTTTTAGGTGTTGGTGGTACTTTATTTGCAGGTGATGTATCTGCACATCGAAGAGCTACTAAAAGCCA |
| 38 | BatLPMO10_SP_USER_rev | ACAACGCCUCTGATAAAACCCTTCATGTTTGTCCTCCTTATTAGTTAATC |
| 39 | Negative_control_USER_fwd | ATGTAAAGAUCCGGCTGTTCAGC |

| | | |
|---|---|---|
| 40 | Negative_control_USER_rev | ATCTTTACATGUTTGTCCTCCTTATTAGTTAATC |
| 41 | LyGo_v1.0.1/2_BB_amyL_USER_rev | AGCGGGGUTTTTTGCGTTAAGCTCTTCGATGCGCC |
| 42 | LyGo_v1.0.1/2_BB_amyQ_USER_rev | AGCGGGGUTTTTTGCGTTAAGCTCTTCGATGGGCTGA |
| 43 | LyGo_V1.0.1/2_BB_aprE_USER_rev | AGCGGGGUTTTTTGCGTTAAGCTCTTCGATGAGCAGCC |
| 44 | LyGo_V1.0.1/2_BB_BatLPMO10_USER_rev | AGCGGGGUTTTTTGCGTTAAGCTCTTCGATGTGCAGATACATCACCTGCAAA |
| 45 | LyGo_V1.0.1_ccdB_USER_fwd | ACCCCGCUTCGGCGGGGTTTTTCGCTACTAAAAGCCAGATAACAGTATGCG |
| 46 | LyGo_V1.0.1_ccdB_USER_rev | AGCCGGATCUTACGAAGAGCTTATATTCCCCAGAACATCAGG |
| 47 | amyQ_mRuby2_fusion_BB_USER_fwd | AGATCCGGCUGTTCAGCG |
| 48 | pBS_CloningSite+terminators_USER_rev2 | AATCGTAAUTATTGGGGACCC |
| | **Sequencing oligonucleotide** | |
| 49 | pBR322_rop_seq | GGTTTTTTCCTGTTTGGTCAC |
| 50 | T7P | TAATACGACTCACTATAGGGGAATTG |
| 51 | T7T | GCTCAGCGGTGGCAGCAGCCAACTCAGCTT |
| 52 | pBS_seq2_fwd | CGCGTCAGGTTAGTGAC |
| 53 | pBS_seq3_rev | TATGAGATAATGCCGACTGT |
| 54 | pBS_seq4_fwd | AGTGAATTTAGGAGGCTTACTTGTC |
| 55 | pBS_seq5_rev | GGCGCGTAAGTGGGATATTT |
| 56 | pBS_seq6_fwd | CGCAATTAATGTGAGTTAGCTCAC |
| 57 | 3P_seqF | CACCCTTGATAGCCTTACTATACC |
| 58 | R0_seqF | GATTAACTAATAAGGAGGACAAAC |
| 59 | pBS_seq7_fwd | CTGAACTAGTCTTGGACTCCTG |
| 60 | pBS_seq8_rev | TCACCAATAAAAAACGCCCGG |

(4) Wendel, S.; Fischer, E. C.; Martínez, V.; Seppälä, S.; Nørholm, M. H. H. A nanobody: GFP

bacterial platform that enables functional enzyme display and easy quantification of display capacity. *Microb. Cell Fact.* **2016**, *15* (1), 1–13.

(5) Radeck, J.; Meyer, D.; Lautenschläger, N.; Mascher, T. *Bacillus* SEVA siblings: A Golden Gate-based toolbox to create personalized integrative vectors for *Bacillus subtilis*. *Sci. Rep.* **2017**, *7* (1), 1–13.
